# Assessing the dynamics of *Mycobacterium bovis* infection in three French badger populations

**DOI:** 10.1101/2023.05.31.543041

**Authors:** Clément Calenge, Ariane Payne, Édouard Réveillaud, Céline Richomme, Sébastien Girard, Stéphanie Desvaux

## Abstract

The Sylvatub system is a national surveillance program established in 2011 in France to monitor infections caused by *Mycobacterium bovis*, the main etiologic agent of bovine tuberculosis, in wild species. This participatory program, involving both national and local stakeholders, allowed us to monitor the progression of the infection in three badger populations in clusters covering between 3222 km^2^ and 7698 km^2^ from 2013 to 2019. In each cluster, badgers were trapped and tested for *M. bovis*. Our first aim was to describe the dynamics of the infection in these clusters. We developed a Bayesian model of prevalence accounting for the spatial structure of the cases, the imperfect and variable sensitivity of the diagnostic tests, and the correlation of the infection status of badgers in the same commune caused by local factors (e.g., social structure and proximity to infected farms). This model revealed that the prevalence increased with time in one cluster (Dordogne/Charentes), decreased in the second cluster (Burgundy), and remained stable in the third cluster (Bearn). In all the clusters, the infection was strongly spatially structured, whereas the mean correlation between the infection status of the animals trapped in the same commune was negligible. Our second aim was to develop indicators for monitoring *M. bovis* infection by stakeholders of the program. We used the model to estimate, in each cluster, (i) the mean prevalence level at mid-period, and (ii) the proportion of the badger population that became infected in one year. We then derived two indicators of these two key quantities from a much simpler regression model, and we showed how these two indicators could be easily used to monitor the infection in the three clusters. We showed with simulations that these two simpler indicators were good approximations of these key quantities.

## Introduction

*Mycobacterium bovis* is a bacterium that can be transmitted to several domestic and wild species, and to humans. It is the main etiologic agent for bovine tuberculosis (bTB), a regulated disease that is still detected in cattle in different European countries. When a farm is infected, different control measures can be applied depending on the country and the specific situation of the farm, including the slaughtering of the herd. France has been officially free of bTB since 2001 (Delavenne, Pandolfi, et al., 2019), as less than 0.1% of cattle herds are infected annually. In certain parts of the country, infection is still regularly detected on cattle farms and in wild species, mainly in wild boars and badgers. The main factor of persistence is cattle-to-cattle transmission through between-herd contact (Marsot et al., 2016; Palisson et al., 2016). However, in some areas, a complex multihost system can explain the circulation of *M. bovis* between different compartments (domestic species, wild species and the environment, Réveillaud et al., 2018); however, even if badgers and wild boars are able to transmit *M. bovis* infection to cattle, these species are not considered long-term maintenance hosts in bTB endemic areas in France (Payne, 2014).

However, due to an increasing number of *M. bovis* cases in wild species, a national surveillance program for *M. bovis* in wildlife named ‘Sylvatub’ was launched in September 2011 (Réveillaud et al., 2018; Rivière et al., 2012). This program aims to detect and monitor *M. bovis* infections in wild species such as wild boar (*Sus scrofa*), red deer (*Cervus elaphus*), roe deer (*Capreolus capreolus*) and European badger (*Meles meles*) populations, by means of both event-based and targeted surveillance strategies. Sylvatub is a participatory monitoring program (*sensu* Danielsen et al., 2003), carried out with the help of local stakeholders such as hunters associations, pest control officers, trapper associations, veterinary associations, livestock health defense associations and epidemiologists (Réveillaud et al., 2018). Briefly, depending on the assessed bTB risk in a given department (French administrative division), three levels of surveillance can be implemented. Level 1 is implemented in a department if no domestic or wild animal has been found to be infected (according to the postmortem examination of hunted or found dead animals). Levels 2 and 3, which are of interest for us in this study, are implemented in departments with sporadic outbreaks in cattle (level 2) and in departments with several outbreaks in cattle and/or cases in wildlife (level 3). In level 3 departments, an at-risk area is defined. This at-risk area is composed of an infected area (communes where the infection has been detected in domestic and/or wild animals – a commune being the smallest French administrative subdivision, with median area of 12 km^2^) and a buffer zone (communes neighboring the infected areas). Trapping is carried out in all the communes of the at-risk area. In level 2 departments, a prospection zone is defined within 2 km from the pastures of infected farms and trapping is restricted to this area (for details, see Réveillaud et al., 2018).

Three main clusters of *M. bovis* infection have been discovered in France during the last 20 years in badger and wild boar populations following an increased prevalence on cattle farms (Delavenne, Pandolfi, et al., 2019) and are being followed up by Sylvatub: Burgundy (initially discovered in wild boar in 2002, and in badgers afterward), Dordogne/Charentes (initially discovered in red deer in 2010, and in wild boar and badgers afterward), and Bearn (initially discovered in wild boar in 2005 and in badgers afterward; Fig 2D). The data collected by this program are used to monitor the spatial extent of the infection as well as its progression within these already infected wild populations, by estimating the prevalence level of the infection in badgers in the different clusters. Since the prevalence is simply the proportion of the population that is infected, it is easily understood by nonspecialist local communities, which is important to keep stakeholders informed and involved in the program. Of course, cattle bTB prevalence and incidence are key parameters to monitor, especially when attempting to maintain the national official bTB-free status; however, because of the multihost system in place, monitoring of cattle prevalence alone cannot capture the complex epidemiological situation of bTB in an area. Therefore, estimating such a parameter in wildlife populations, that are easily comparable from one year to another, would also be essential for monitoring the epidemiological situation and evaluating the impact of control measures.

However, an ongoing issue in wildlife epidemiology is the difficulty in estimating prevalence in wild populations, as the sampling of animals used for this estimation cannot be entirely controlled. Indeed, the population is usually sampled using capture methods (e.g., traps for badgers), and the prevalence is usually estimated from the sample of captured animals, under the assumption that these animals are a random sample of the population (which therefore ignores possible capture bias such as the uneven behavior of the animals toward the traps and the logistical constraints that can affect the placement of traps). Moreover, in the case of participatory monitoring programs, the participating local communities generally already have their own objectives (e.g., wanting to trap more animals close to some given farms during certain years, and close to others during other years) in addition to the Sylvatub objectives. Thus, monitoring protocols cannot be too rigid in participatory programs involving volunteers (e.g., Pocock et al., 2015). However, the spatial structure of the infection must be considered when estimating the prevalence or any related indicator in a given population.

In addition, another estimation problem occurs when the sampled species is characterized by a social structure that makes trapped animals nonindependent from each other. For example, badgers typically live in social groups that share the same sett and mutually defend a group territory (Roper, 2010). As a consequence, a correlation of infection status is expected among animals trapped at a given place (e.g., Delahay et al., 2000): when one trapped animal is infected, it is likely that other animals trapped at the same place belong to the same group, and are therefore also infected. Moreover, it has been shown that bTB infections in badgers and cattle are spatially associated (Bouchez-Zacria, Courcoul, et al., 2018; Bouchez-Zacria, Payne, et al., 2023); therefore badgers trapped near an infected farm are more likely to be infected. Not accounting for this correlation when estimating the prevalence may lead to an overestimation of precision (Hisakado et al., 2006).

A final difficulty occurs when the sensitivity and specificity of the tests used for diagnosis are not perfect: not all infected (resp. non-infected) animals are identified as such by these tests; there may be false positives and false negatives. Ignoring this imperfect measure of infection status can lead to biased estimation of the prevalence (Dohoo et al., 2003). Moreover, if the tests used for this diagnosis (and the corresponding sensitivity/specificity) change with time, the assessment of infection progression based on the uncorrected prevalence estimation may also be biased.

In this study, we focused on the targeted surveillance of badgers, which was carried out in communes characterized by surveillance levels 2 and 3 (representing 80% of the data collected in the framework of Sylvatub between 2013 and 2019): in each identified bTB cluster, traps were set up by members of Sylvatub in the communes of the at-risk areas, and *M. bovis* infection was sought in a subsample of the trapped badgers (the proportion and spatial distribution of tested animals depend on the number of trapped animals, trap location and annual sampling objectives). We used these trapping data to develop a complex Bayesian model and provide insight into how the proportion of infected badgers varied in space and time in the three French bTB clusters, accounting for the complex spatial structure of the infection, the correlation between the infection status of animals trapped at the same place and the limited sensitivity of the diagnostic tests. Then, we used this model as a basis for developing simpler indicators of the prevalence that also account for all the aforementioned difficulties. These simpler indicators can be easily understood by all stakeholders and used to monitor both the mean prevalence level and the mean prevalence trend over a given period. The work in this paper is summarized in Fig 1.

**Figure 1.**
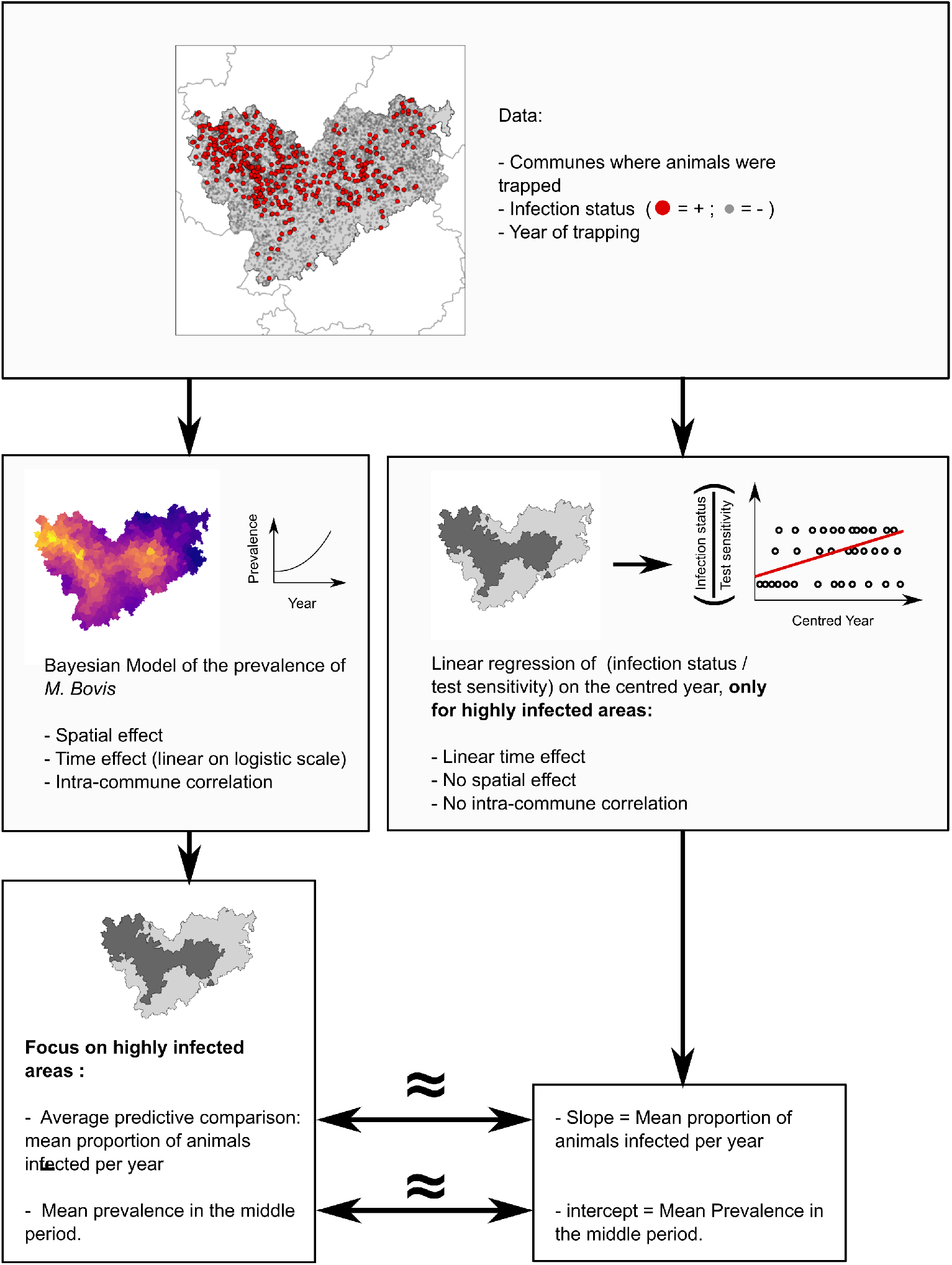
Summary of the models fitted in this paper. For each cluster (illustrated herein with the Dordogne/Charentes cluster), our dataset consisted of a sample of badgers trapped in different communes during different years, and tested for *M. bovis*. We first fit a complex Bayesian model to this dataset accounting for many characteristics of the infection (left). We then focused on highly infected communes and used the average predictive comparisons to estimate the mean proportion of the cluster population infected in one year. Additionally, we used the model to estimate the mean prevalence of these infected communes during the year in the middle of the study period. We then fit a much simpler linear regression (right) on the data collected from the highly infected communes, which allowed us to directly estimate the mean proportion of the cluster population becoming infected in one year, and the mean prevalence in the cluster population during the year in the middle of the study period. The simulations indicate that the two approaches returned nearly identical results.

## Material and methods

### Sylvatub program and database

The national surveillance system is described in detail in Réveillaud et al. (2018). Briefly, in the communes from level 2 and 3 departments (i.e., communes from infected areas), trapping and culling badgers is implemented as a control measure to reduce badger abundance. To do so, licensed field stakeholders (hunters, trappers, pest control officers) trap badgers, mostly between March and August. The regulatory guidelines for badger trapping are uniform across the three clusters, and are defined by the French Ministry of Ecology and Sustainable Development, through the ministerial order of January 29th, 2007. Only two types of traps are authorized in France: stop snare (i.e., snare with a mechanism that stops the noose from closing too tightly) and cage traps. Night shooting is also an option in level 3 communes. Nevertheless, with a few exceptions, French trappers predominantly utilize stop snares. Given the participatory nature of the Sylvatub program, local trappers retain the autonomy to decide on the number of traps, trap nights, and their placement. However, the Sylvatub program encourages however trapping near infected farms (technical directive from the French Ministry of Agriculture DGAL/SDSPA/2018-708).

The trapped badgers are culled, and sent to the local veterinary laboratory for necropsy and *M. bovis* testing following the framework of Sylvatub. Not all dead animals are tested; the national guidelines are to analyze at most two animals in each commune and each year. The choice of analyzed animals among trapped badgers was left to the local partners of the network. Given that the infection status of trapped badgers is seldom discernible from external observations (as most TB lesions diagnosed in badgers are internal, as noted by Réveillaud et al. (2018)), we are confident that there was no sampling bias directly related to the infection status of animals. While trapping efforts were intensified near infected farms to control the density of badgers in proximity to these areas, Sylvatub guidelines encouraged the analysis of badgers distributed spatially as uniformly as possible. In practice, we observed that the badgers of a commune sent for analysis were often the first two trapped badgers.

In the following, we assume that the animals sent to the laboratory are a random sample of the trapped animals. This guideline to test at most two animals was intended more as a general recommendation rather than a strict fieldwork requirement. Although fieldworkers generally adhered to these guidelines during the study period, approximately 25% of the communes trapped and analyzed more than two animals per year on average. In all three clusters, between 12% and 13% of the communes trapped and analyzed more than 21 badgers over the 7-year period, and between 3 and 4% of the communes trapped more than 36 badgers during this period. Note that our statistical approach assumed the ignorability of the sampling; in other words, we assumed that infected and noninfected badgers are characterized by equal trappabilities. The results of the test for each analyzed animal are stored in a local database and subsequently compiled in the national Sylvatub database.

As Sylvatub was launched in 2011 and was not yet well-established before 2013, our study period therefore covered 2013 to 2019. The set of communes where targeted surveillance was authorized for at least one year between 2013 and 2019 was used to define three main spatially connected sets, which are hereafter called *M. bovis* clusters (Fig 2D). The Dordogne/Charentes cluster covers 7698 km^2^ and is composed of 413 communes; the Burgundy cluster covers 4254 km^2^ and is composed of 254 communes; and the Bearn cluster covers 3222 km^2^ and is composed of 196 communes. The median surface area of a commune is 12 km^2^ (interquartile range: 7.2 km^2^ to 18.3 km^2^). Note that we lack precise information regarding the social group size of badgers in the three clusters. Jacquier et al. (2021) employed a standard methodology, utilizing camera traps and genetic identification, to estimate badger density across multiple sites in France, including the three clusters of interest. These authors showed that the badger density was highest in the Dordogne/Charentes cluster (6.18 badgers / km^2^), followed by the Bearn cluster (5.39 badgers / km^2^), and the Burgundy cluster (two sites were studied by these authors in this cluster and were characterized by a density of 4.08 and 4.22 badgers / km^2^). For comparison, the mean density across the 13 sites studied by these authors, distributed across the entire metropolitan region of France was 5.85 badgers / km^2^ – SD = 3.25 badgers / km^2^).

**Figure 2.**
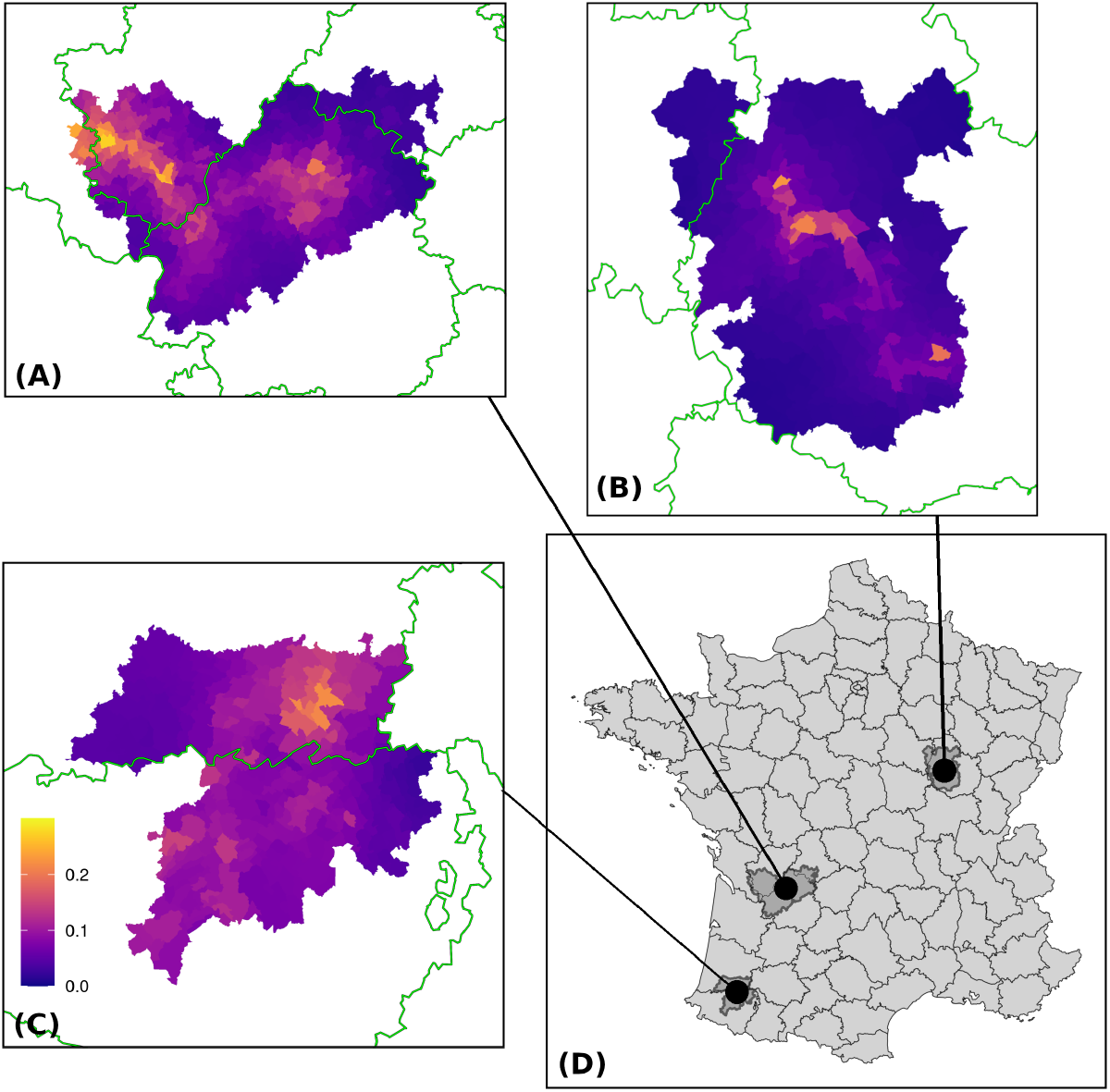
Location of the three *M. bovis* clusters in France (D). The boundaries of the French departments are displayed on this map, as well as the median prevalence estimated by our Bayesian model for each commune in the Dordogne/Charentes cluster (A), the Burgundy cluster (B), and the Bearn cluster (C). A common color scale is used for all clusters (inset in the Bearn map).

Following necropsy, two types of first-line tests were carried out on the animal samples. Pools of lymph nodes (retropharyngeal, pulmonary and mesenteric) and organs with gross lesions were used in the analysis. The type of analysis depended on the period: (i) from 2013 to 2015, the first-line test was the bacterial culture performed on the sampled tissues of all tested animals, following the protocol established by the French NRL (NFU 47-104) for the isolation of *M. bovis*; (ii) since 2016, the first-line test has been real-line PCR performed after DNA extraction from the sampled tissues. Molecular typing (spoligotyping) was performed either on Mycobacterium tuberculosis complex (MTBC) isolates or directly on PCR-positive sample DNA (see Réveillaud et al., 2018, for technical details on these two procedures). The sensitivities of the two techniques differ: the sensitivity of microbiological cultures is estimated to be 50%, whereas the sensitivity of the PCR is estimated to be 75% (Réveillaud et al., 2018; Riviere et al., 2015). The specificity should be equal to 100% for these two tests (i.e., no false positives).

During the study period, 4590 badgers were trapped and sent to the laboratory in Dordogne/Charentes, among which 4379 badgers were actually tested. Interpretable results were obtained for 4323 of them (i.e., on average 1.5 badgers per commune and per year; interquartile range: 0 animals to 2 animals tested per commune and per year). In Burgundy, 3042 badgers were trapped and sent to the laboratory, among whom 2900 were actually tested; interpretable results were obtained for 2786 of them (on average 1.56 animals were tested per commune and per year; interquartile range: 0 to 2 animals were tested per commune and per year). Finally, in Bearn, 2223 badgers were trapped and sent, among which 1999 were tested; interpretable results were obtained for 1970 (on average 1.43 animals were tested per commune and per year; interquartile range: 0 to 2 animals were tested per commune and per year).

For each trapped animal, the following data were stored: date of trapping, name of the commune where the animal was trapped, results of the test (*M. bovis* positive, *M. bovis* negative), type of first-line test carried out (bacterial culture; PCR), date of the analyses, surveillance level of the commune of trapping, and sex and age class (young; adult) of the animals (although this latter information is not systematically reported by the field partners).

### A Bayesian model of infection

#### Model fit

For each of the three *M. bovis* clusters, we fitted a Bayesian model describing the dynamics of the infection process. Consider one particular cluster. Let *N*_*it*_ be the known number of badgers trapped and tested in commune *i* during year *t*. Let *y*_*it*_ be the unknown number of badgers actually infected among those *N*_*it*_ animals. Let *z*_*it*_ be the known number of animals for which the test indicated *M. bovis* infection among those *y*_*it*_ infected animals. We fitted the following hierarchical Bayesian model:

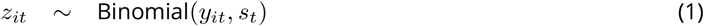

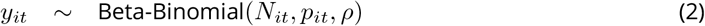

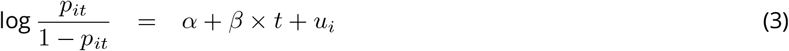

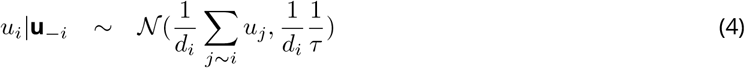

Equation (1) accounts for the known sensitivity *s*_*t*_ of the tests used during year *t* (i.e., *s*_*t*_ = 0.5 for microbiological culture, and *s*_*t*_ = 0.75 for PCR): the number *z*_*it*_ of animals for which an *M. bovis* infection was diagnosed is a random subset of the unknown number *y*_*it*_ of animals actually infected (which is a latent variable in this model). Each infected animal is detected as such with a known probability *s*_*t*_.

We assumed a beta-binomial distribution for the unknown number of infected animals *y*_*it*_ (Equation (2)). This distribution accounts for a possible correlation between the infection status of the animals trapped in the same year in the same commune, and is parameterized by the known number *N*_*it*_ of badgers trapped in commune *i* during year *t*, the unknown prevalence *p*_*it*_ of *M. bovis* infection in commune *i* and year *t*, and the unknown correlation coefficient *ρ* (estimated by the model fit) between the infection status of the animals trapped in the same commune. The parameterization of the beta-binomial distribution as a function of a probability (here, the prevalence) and a correlation coefficient was proposed by Hisakado et al. (2006) as a means to account for the correlation between binary variables in binomial counts. Appendix A gives the formal expression of this distribution with this parameterization, and discusses how it relates to the parameterization classically used by statistical software such as R.

The prevalence *p*_*it*_ is itself modeled by a logistic regression depending on a commune effect and a linear year effect (also estimated by the model fit; Equation (3)). The effects *u*_*i*_ of the communes on the prevalence are not independent of each other. Indeed, because of the strong spatial structure of the infection in the clusters, there is a high probability that the prevalence of infection is high in a commune if it is high in neighboring communes. We account for this spatial autocorrelation of the commune effects by modeling these random effects *u*_*i*_ with an intrinsic conditional autoregressive (iCAR) model (Equation (4), see also Rue and Held, 2005). Thus, the random effect *u*_*i*_ of commune *i* is assumed to be drawn from a Gaussian distribution with a mean equal to the mean of the random effects of neighboring communes. In Equation (4), *i ∼ j* means that commune *i* shares a boundary with commune *j*, **u***−i* is the vector of commune random effects excluding the effect *u*_*i*_, and *d*_*i*_ is the number of communes sharing a boundary with commune *i*. The parameter *τ* is estimated by the model fit, which describes the precision (inverse of the variance) of the random effects *u*_*i*_.

We defined weakly informative priors for the parameters of the model. We fitted this model by Markov chain Monte Carlo (MCMC) using 4 chains of 1 million iterations each after a burn-in period of 3000 iterations. To save some memory space, we thinned the chains by selecting one sample every 1000 iterations. We checked the mixing properties of the chains both visually and using the diagnostic method of Gelman and Rubin (1992). We checked the goodness of fit of our model using the approach recommended by Gelman and Meng (1996): each MCMC iteration *r* generated a sampled value *θ*^*r*^ of the vector of parameters of interest *θ* = (*τ, α, β, ρ*, **u**)^*t*^. For each simulated value *θ*^*r*^, we simulated a replication of the Sylvatub dataset (i.e., we simulated a random infection status for each trapped animal of the dataset with the fitted model parameterized by the vector simulated by the *r*-th MCMC iteration). We then compared summary statistics calculated on the observed Sylvatub dataset with the distribution of the same statistics derived from the simulated datasets. All these checks indicated a satisfactory fit of the model (see Appendix D for details on these checks and on model fit).

### Estimation of the prevalence level and trend from the model

First, we used the fitted model to estimate the trend over time of the prevalence in each cluster. On a logit scale, the average change in the prevalence with time is reflected by the coefficient *β* in Equation (3). It is well known that in a logistic regression, the exponential of a coefficient (here *β*) is equal to the odds ratio of the corresponding variable (here the time *t*), i.e., *{p*_*t*_/(1 *− p*_*t*_)*}*/*{p*_*t−*1_/(1 *− p*_*t−*1_)*}*, which in our model measures the amount by which the odds *p*/(1 *− p*) of the infection are multiplied in one year (Hosmer and Lemeshow, 2000, p. 50). However, odds ratios are difficult for stakeholders to understand, which can be problematic in a participatory program context. As noted by King and Zeng (2002), “*we have found no author who claims to be more comfortable communicating with the general public using an odds ratio*. Similarly, Gelman and Hill (2006, p. 83) reported that “*we find that the concept of odds can be somewhat difficult to understand, and odds ratios are even more obscure. Therefore, we prefer to interpret coefficients on the original scale of the data*”. In this section, we follow this last recommendation, by calculating the average rate of change in the prevalence in a cluster using the fitted model.

Due to both the nonlinearity of the logit transform used in the model and the strong spatialization of the infection, estimating from the model of the average proportion of animals becoming infected in one year can be difficult. Gelman and Pardoe (2007) proposed an approach to estimate this rate of change, based on the concept of predictive comparison. For a given commune *v* and a given value of the vector of parameters *θ* of the model, the predictive comparison measures the expected rate of change in the prevalence *p* when the year changes from *t*^(1)^ to *t*^(2)^:

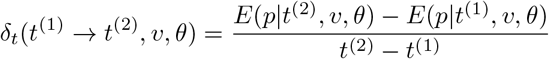

This quantity, easily calculated with our model, varies as a function of these inputs (the years compared, the commune, and the value of the vector of parameters). To summarize the overall effect of the year on the prevalence in a given dataset, Gelman and Pardoe (2007) proposed calculating the mean value Δ_*t*_ of the predicted comparisons over the probability distribution of the inputs (years and communes) estimated with the data, and over the posterior distribution of the parameters. This averaging is equivalent to considering all pairs of animals (*i, j*) in the data, corresponding to pairs of transitions of (*t*_*i*_, *v*_*i*_) to (*t*_*j*_, *v*_*j*_) in which the commune *v*_*i*_ = *v*_*j*_ is held constant. Technical details on the calculation of the average predictive comparison (APC) are given in Appendix B. When positive, the APC estimated the proportion of the animals that became infected within one year in each cluster. Conversely, negative values of the APC indicated a decrease in the prevalence within one year. This reduction results from a combination of factors, including the death of infected animals, the birth of uninfected animals, or a decrease in the infection rate.

The APC provides an index of the dynamics of the infection in a cluster. We also estimated another statistic summarizing the mean prevalence level in a cluster during the study period. Because the prevalence varied spatially and temporally, we used Equation (3) to estimate the expected prevalence in each commune during the middle year of our study period (i.e., for the year *t* =2016), and we averaged it over the communes of the cluster. This gave an idea of the importance of the infection in each cluster during the study period.

The calculation of both the APC and the mean prevalence level during the middle year was restricted to the set of highly infected communes (i.e., communes for which *u*_*i*_ *>* 0 in Equation (4)) to allow comparison with the simpler indicators developed in the next section.

### Development of simple indicators of ***M. bovis*** prevalence level and trend

Although the model developed in the previous sections is useful for understanding the spatialization and dynamics of the infection process, it is too complex to be used on a regular basis by the stakeholders of Sylvatub to assess how the prevalence level changes with time. Instead, in this section, we propose two new indicators that can be estimated with the trapping data collected by the network. These indicators estimate in a simpler way the same quantities that were introduced in the last subsection, i.e., the mean prevalence level in the middle year of the study period and the mean proportion of animals becoming infected in one year.

Consider a given *M. bovis* cluster during a given study period of several years *t* = 1, 2, …, *T*, during which *n* animals were collected via the Sylvatub network. For each animal *i*, let *B*_*i*_ be the infection status returned by the test (coded as 0/1) and *s*_*i*_ be the sensitivity of the test used for this diagnosis. We can derive two useful indicators with classical simple linear regression fitted to the set of animals trapped during the study period:

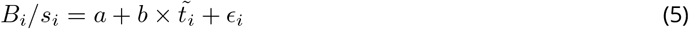

Where 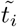 is the centred year (i.e. 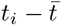, where 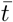 is the middle year of the study period), and *ϵ*_*i*_ is a residual. In this model, the coefficient *a* corresponds to the average prevalence observed in the middle year of the study period, and the coefficient *b* corresponds to the proportion of the badger population that becomes infected during a year on average during the study period (i.e., the same quantity as the APC calculated for the Bayesian model; see Appendix C for a detailed explanation of the rationale).

This approach accounts for the imperfect sensitivity of the tests used for *M. bovis* diagnosis, but does not account for the spatial structure of the infection in the cluster under study. We show (see the Results section) that there is a very strong spatial structure of the infection in the three *M. bovis* clusters. Therefore, not accounting for this structure can lead to biased estimates if the trapping pressure in highly infected areas varies between years. We therefore suggest calculating these prevalence indicators by focusing only on highly infected communes (i.e. communes characterized by an estimated random effect *u*_*i*_ greater than 0 in Equation (4)), so that the remaining unaccounted spatial variability of infection can be ignored. This approach also ignores the correlation possibly caused by the social structure of the badger population and by other local factors (e.g. proximity to an infected farm); however, we show that this correlation is negligible in the three clusters (see Results).

### Assessing the indicators with the Bayesian model

The complex Bayesian model described by equations (1) to (4) and the simpler regression model described by Equation (5) are two models of the same process, though the latter is much simpler. Both can be used to estimate the mean prevalence level during the middle year of the study period and the mean proportion of the population becoming infected in one year in a cluster. The simple regression model imperfectly accounts for the spatial structure of the infection and ignores the correlation caused by local factors (e.g. social structure, proximity to infected farms); however, this approach is much easier for stakeholders to understand and implement. This latter model is therefore proposed for stakeholders as a means to monitor infection in a cluster.

We carried out three sets of simulations to assess the ability of the simpler regression model to estimate the two target quantities. The first set was designed to assess the ability of our regression model to estimate the trend over time of the prevalence in various situations that might be encountered in reality (that is, either an initially rare but increasing infection or an already widespread infection with different trends). The second set was designed to assess the ability of our regression model to estimate the mean prevalence level of the infection at a variety of actual levels (from rare to very frequent infection). The last set was designed to assess the robustness of our approach to violations of the hypotheses on which it relies (strong spatial heterogeneity remaining even in highly infected communes, spatial structure of the infection changing with time, nonrandom sampling).

For all these simulations, we used the Dordogne/Charentes cluster as an example. In all the cases, we simulated datasets covering 7 years in this cluster. We generated a sample of trapped animals for each year and for each commune *i* of the cluster from a binomial negative distribution with mean *μ*_*i*_ and dispersion parameter *θ* = 0.48 (this value was estimated from our dataset by maximum likelihood); in the first and second sets of simulations, four possible values of *μ*_*i*_ = *μ*, corresponding to four levels of trapping pressure, were simulated: *μ* = 0.5 animals trapped per commune and per year on average, and *μ* = 1, *μ* = 3 and *μ* = 10 (as a point of comparison, remember that in our dataset, *μ ≈* 1.5 in all clusters). In the third set of simulations, *μ*_*i*_ varied among communes (see below). For each simulated animal, we simulated the probability of infection with the help of Equation (3). Different values of the slope *β* and intercept *α* were specified for the different simulation situations (see below). We simulated an iCAR process to generate random commune effects *u*_*i*_ using Equation (4). We set *τ* = 0.73 for this process in the first and second sets of simulations (corresponding to the mean value estimated by the model with the Sylvatub dataset in the Dordogne/Charentes Cluster, see Results). We used another value of *τ* for the third set of simulations (see below). For each animal, we calculated the probability of infection *p*_*it*_ from the vector (*α, β, {u*_*i*_*}*) with Equation (3). We then simulated the random infection status of each animal using Equation (2), fixing the correlation coefficient *ρ* = 0.04 (which also corresponded to the value estimated in the Dordogne/Charentes cluster using the Sylvatub dataset, see Results). Finally, we used Equation (1) to simulate an imperfect but variable sensitivity (sensitivity equal to 0.5 during the first three years and 0.75 during the last four years).

In the first set of simulations, we wanted to assess the ability of our regression model to estimate the trend over time of the prevalence in two different situations with regard to its change with time: (i) a rare infection that becomes more widespread with time: we simulated an *M. bovis* infection rarely present in the study area during the first year (*≈* 5% of the animals were infected in a typical commune of the cluster), with the prevalence increasing with time. More precisely, we set the intercept *α* = *−*3.1 in Model (3) and the slope *β* of the year was randomly drawn from a uniform distribution bounded between 0 and 0.4; (ii) an already widespread infection with different trends: we simulated frequent infection during the first year of the study period (*≈* 20% of the animals were infected in a typical commune) with an either increasing or decreasing prevalence. More precisely, we set the intercept *α* = *−*1.38 and the slope *β* randomly drawn from a uniform distribution bounded between -0.4 and 0.4. For each combination of trapping pressure *μ* and simulated situation (either low but increasing prevalence or high prevalence), we simulated 1000 datasets. For each dataset, we estimated the true proportion Δ_*u*_ of animals in the area that became infected within one year in the highly infected communes (i.e. those with simulated random effects greater than 0) via the APC procedure. We applied linear regression (5) to the data simulated in these communes. We then compared the estimated slope *b* with the APC Δ_*u*_ of the simulated model, which should in theory be equal if the two models are equivalent.

In the second set of simulations, we wanted to assess the ability of our regression model to estimate the mean prevalence level during the middle year. We simulated the data with our Bayesian model, using different values of the intercept *α* = *−*4, *−*3, *−*2, *−*1, 0, which describes different mean prevalence levels. We then randomly sampled a slope *β* from a uniform distribution bounded between -0.4 and 0.4. We simulated 1000 datasets for each combination of intercept *α* and trapping pressure *μ*. For each simulated dataset, we considered only the highly infected communes (i.e., those with *u*_*i*_ *>* 0) and we calculated the true mean prevalence over the area during the middle year of the study period. We then applied linear regression (5) to the data simulated in these communes. We then compared the estimated slope *a* with this true mean prevalence, which should be equal if the two models are equivalent.

In the third set of simulations, we aimed to assess the robustness of our model to the violation of two underlying hypotheses: (i) ignorability of the remaining spatial structure of the prevalence when the regression model is applied only to the data coming from highly infected communes and (ii) additivity of the space and time effects on the prevalence. In these two situations, we simulated the data with our Bayesian model using two different values of the intercept *α* = *−*2, 0, representing different mean prevalence levels. We then randomly sampled a slope *β* in a uniform distribution bounded between -0.4 and 0.4. To test the effect of the violation of the first hypothesis, we simulated random commune effects *u*_*i*_ using Equation (4), setting a very low value *τ* = 0.1, corresponding to very strong spatial heterogeneity. To test the effects of violating the second hypothesis (additivity of space and time effects), we simulated the spatial structure of the infection changing with time. More precisely, we simulated two sets of commune effects,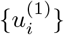 and 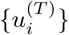, describing the spatial structure at the start and end of the study period, respectively (using *τ* = 0.73 in both cases). The set of random effects used at time *t* was calculated by 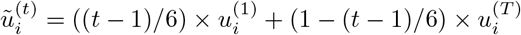 . In the two tested situations, we estimated the two parameters of interest (intercept and slope of the regression model) and compared them to the theoretical values used for simulation. In this third set of simulations, two sampling schemes were compared to demonstrate how directed sampling can exacerbate the effect of the violation of underlying hypotheses: random sampling with *μ*_*i*_ = 2 and directed sampling where the mean number of animals in a commune was proportional to the mean prevalence in the commune, i.e. *μ*_*i*_ = 2 *× M ×* exp(*u*_*i*_)/(Σ_*j*_ exp(*u*_*j*_)) (where *M* is the number of communes).

### Computational aspects

All our analyses and simulations were carried out with R software (R Core Team, 2023). We used the package nimble for model fit (Valpine et al., 2017), coda for the analysis of the fit (Plummer et al., 2006), and tidyverse (Wickham and Grolemund, 2017) and `ggplot2` (Wickham, 2016) for data manipulation and graphical display. We programmed an R package named `badgertub`, available at https://github.com/ClementCalenge/badgertub, containing all the code and data used to fit the model. The package can be installed in R with the package devtools using the function `devtools::install_github(“ClementCalenge/badgertub”, ref=“main”)`. This package includes a vignette describing how the user can easily reproduce the model fit and simulations (vignette available with the command `vignette(“badgertub”)` once the package has been installed). This vignette also serves as the supplementary material for the paper and contains additional information on our model (e.g., precision on the parameterization of the beta-binomial distribution, and a formal description of the iCAR model).

## Results

### Collected data and model fit

The number of animals trapped in each *M. bovis* cluster during each year of the study period is presented in Tab 1, as is the proportion of these animals diagnosed as infected with *M. bovis*. Note that even though it is challenging to interpret the observed temporal changes in prevalence (as this proportion does not account for all the factors that influence the prevalence, i.e., inhomogeneous prevalence patterns in space, sensitivity of the tests used increasing with time, etc.), this table clearly demonstrates the overall temporal change observed in each cluster, i.e. a strong increase in Dordogne/Charentes, a decrease in Burgundy, and a moderate increase in Bearn.

**Table 1.** Number *N* of trapped badgers during each year of the study period in each one of the three *M. bovis* clusters, and proportion *p* of the badgers that were diagnosed as infected with *M. bovis* (the corresponding number *n* of infected animals is in parentheses).

| Year | Dordogne/Charentes |  | Burgundy |  | Bearn |  |
| --- | --- | --- | --- | --- | --- | --- |
|  | N | p (n) | N | p (n) | N | p (n) |
| 2013 | 449 | 0.024 (11) | 310 | 0.094 (29) | 381 | 0.045 (17) |
| 2014 | 637 | 0.036 (23) | 636 | 0.05 (32) | 387 | 0.047 (18) |
| 2015 | 787 | 0.051 (40) | 479 | 0.044 (21) | 189 | 0.042 (8) |
| 2016 | 639 | 0.036 (23) | 429 | 0.037 (16) | 287 | 0.077 (22) |
| 2017 | 700 | 0.056 (39) | 310 | 0.019 (6) | 320 | 0.053 (17) |
| 2018 | 502 | 0.1 (50) | 303 | 0.003 (1) | 204 | 0.059 (12) |
| 2019 | 609 | 0.1 (61) | 319 | 0.041 (13) | 202 | 0.109 (22) |

The model provides a clearer point of view on the infection process. The estimated parameters of the model for each cluster are presented in Tab 2. The abundance of data available in the three clusters results in precise estimations, as evident from the narrow width of the credible intervals for all parameters in this table. The situation was contrasted among the three clusters: the infection strongly decreased in Burgundy, strongly increased in Charentes/Dordogne and seemed stable in Bearn, as revealed by both the slope *β* of the year in the model and the APC (i.e., the proportion of animals becoming infected in one year). The correlation *ρ* between the infection status of animals trapped in the same commune was actually rather small in all the clusters (*≈* 0.03), revealing that the correlation caused by local factors (social structure, local environment, etc.) did not cause a strong dependency between animals in a commune. Conversely, a strong spatial structure was evident in all three studied clusters, revealing distinct patterns of highly infected areas and low-risk areas within each cluster (see Fig 2). Specifically, the set of highly infected communes formed a connected subset of communes (i.e. a unique subarea) in the three clusters, except the Dordogne-Charentes cluster, where two highly infected communes were located only a few kilometers away from the main subarea. Furthermore, the proportion of trapped animals diagnosed as infected was greater in the highly infected communes than in the other communes (focusing on 2017–2019 to limit temporal changes: 16% in highly infected communes of Dordogne-Charentes vs. 3% in other communes; 11% in highly infected communes of Bearn vs. 0.75% in other communes; and focusing on 2013–2015 in Burgundy, when the infection rate was still noteworthy; 10.6% in the highly infected communes vs. 0% in the other communes).

In the three clusters, there was close agreement between the parameters estimated by the Bayesian model and the same parameters estimated by simple linear regression (Tab 2), although the mean prevalence seemed to be slightly overestimated by the regression approach in the three clusters or slightly underestimated by the Bayesian model.

**Table 2.** Main results derived from the model fit to the three *M. bovis* clusters. We present here: (i) the parameters of interest in the model (the first three rows are the parameters of the model: slope *β* associated with the year, correlation coefficient *ρ* between the infection status of animals trapped in the same commune, and standard deviation 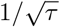 of the commune effects); (ii) the average predictive comparison (APC) estimating the proportion of the population becoming infected in one year as estimated by the complex Bayesian model and by the simpler regression in the highly infected communes (see text); (iii) the mean prevalence level in the highly infected communes in the middle year of the study period (see text) as estimated by the complex Bayesian model and by the regression. For each parameter and each cluster, we give the point estimate (mean of the posterior distribution) and the 90% credible interval.

| Parameter | Dordogne/Charentes | Burgundy | Bearn |
| --- | --- | --- | --- |
| $\beta$ | 0.18 [0.11, 0.25] | -0.29 [-0.39, -0.2] | 0.05 [-0.04, 0.14] |
| $\rho$ | 0.04 [0.02, 0.08] | 0.02 [0.01, 0.04] | 0.04 [0.02, 0.08] |
| $1/\sqrt{\tau}$ | 1.17 [0.87, 1.5] | 1.58 [1.11, 2.04] | 1.08 [0.69, 1.54] |
| APC (model) | 0.018 [0.009, 0.027] | -0.028 [-0.039, -0.017] | 0.005 [-0.008, 0.018] |
| APC (regression) | 0.029 [0.02, 0.037] | -0.034 [-0.043, -0.024] | 0.008 [-0.002, 0.018] |
| Mean Prevalence (model) | 0.126 [0.109, 0.143] | 0.08 [0.065, 0.097] | 0.112 [0.092, 0.134] |
| Mean Prevalence (regression) | 0.157 [0.141, 0.174] | 0.117 [0.098, 0.136] | 0.133 [0.113, 0.153] |

The first two sets of simulations revealed that the two indicators correctly estimated the mean prevalence and the mean proportion of animals becoming infected fixed in our simulated situations. On the one hand, the first set of simulations of two contrasting situations (high prevalence or low and increasing prevalence) showed that the slope of the year in the regression agreed with the true simulated proportion of animals becoming infected in one year, regardless of the simulated trapping pressure (Fig 3). Of course, the uncertainty was greater when the trapping pressure was lower (the cloud of points was more dispersed around the line *y* = *x* when *μ* was low); however, this indicator correctly estimated the target proportion.

**Figure 3.**
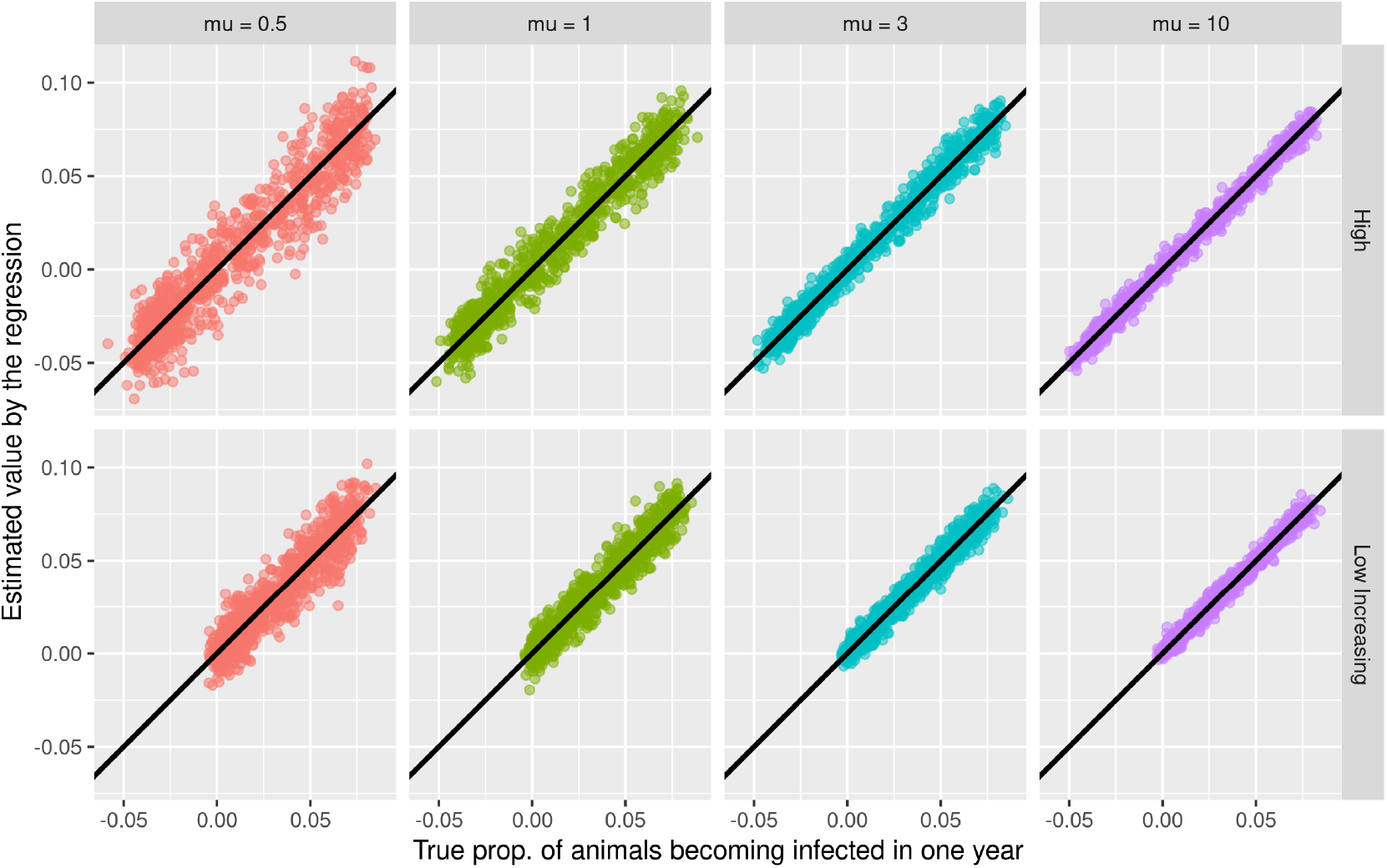
Comparison of the proportion of animals infected in one year estimated using the regression indicator (see text) with the true value, estimated by simulations for two different situations (high prevalence = top row; low but increasing prevalence = bottom row) and different trapping pressures (mu corresponds to the mean number of animals trapped per commune). The straight line is the line of equation *y* = *x*.

On the other hand, the second set of simulations of different prevalence levels under different trapping pressures showed that there was close agreement between the intercept of the linear regression and the true mean prevalence level during the middle year in highly infected communes (Fig 4). Similarly, the uncertainty was greater for low trapping pressures.

Since we used linear regression to estimate our two indicators, we derived confidence intervals for these two parameters using the classical formulas derived from the normal theory. We calculated the coverage probability of the 95% confidence intervals for the different simulated situations (Tab 3 and Tab 4). In these first two sets of simulations, the coverage probabilities of the 95% confidence intervals for the two indicators were closer to 90% than to 95% for moderate trapping pressure. When the trapping pressure was extremely high (i.e., 10 animals trapped on average in each commune of a cluster), the coverage probability of the 95% confidence interval decreased to *≈* 80% for the proportion of animals becoming infected in one year and to *≈* 60% for the mean prevalence level during the middle year.

**Figure 4.**
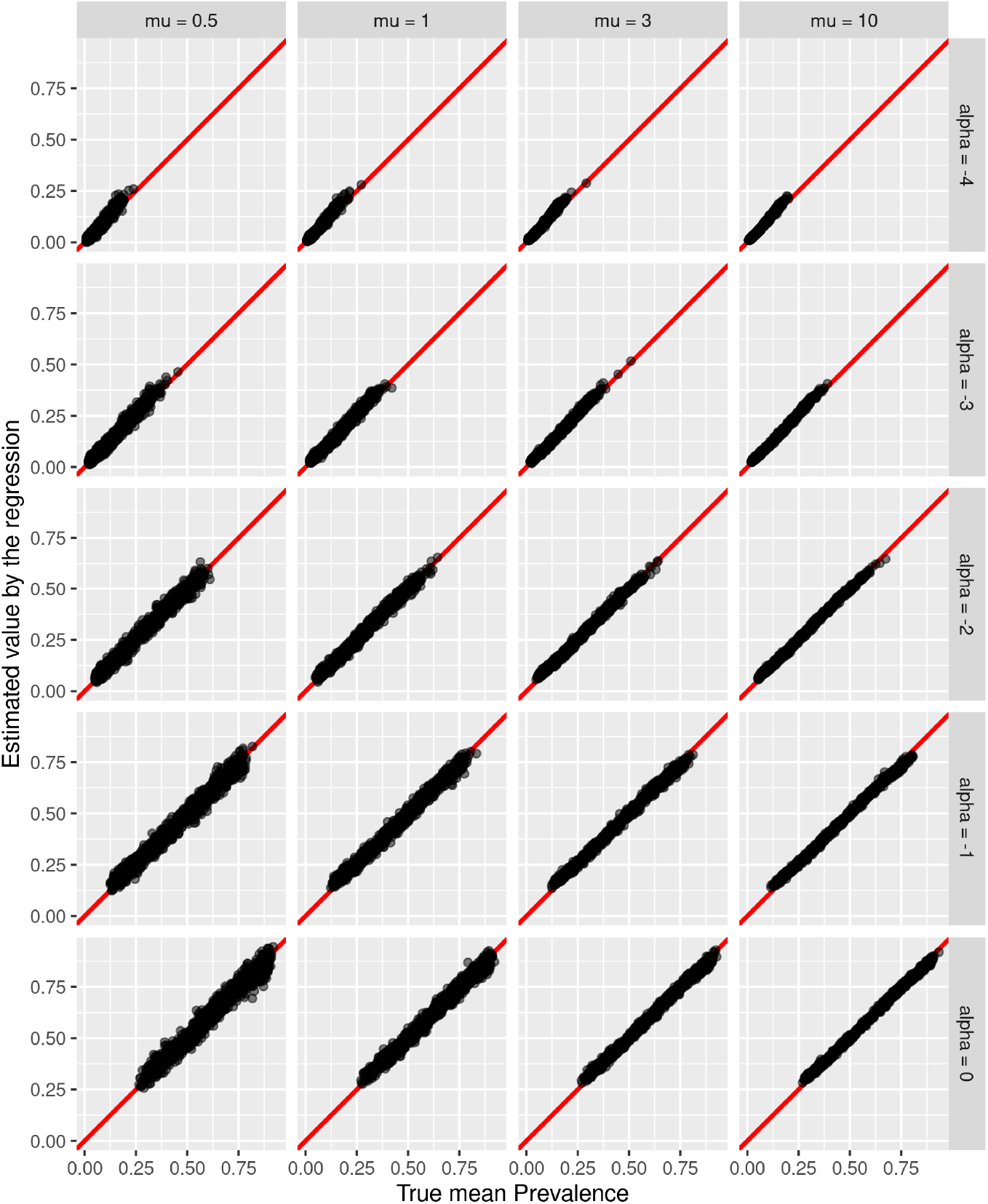
Comparison of the mean prevalence level during the middle year estimated using the regression indicator (see text) with the true value, estimated by simulations for five different prevalence levels (simulated by fixing different values of the intercept alpha) and the different trapping pressures (mu corresponds to the mean number of animals trapped per commune). The straight line is the line of equation *y* = *x*.

**Table 3.** Coverage probability of the 95% confidence interval on the proportion of animals in a cluster infected in one year estimated with simple linear regression, estimated by simulations for the two tested settings (either high prevalence or low but increasing prevalence) and the 4 trapping pressures. The value of *μ* corresponds to the mean number of animals trapped in each commune.

| Situation | Trapping Pressure | Coverage Probability |
| --- | --- | --- |
| High | $\mu = 0.5$ | 0.93 |
| High | $\mu = 1$ | 0.94 |
| High | $\mu = 3$ | 0.91 |
| High | $\mu = 10$ | 0.84 |
| Low Increasing | $\mu = 0.5$ | 0.94 |
| Low Increasing | $\mu = 1$ | 0.94 |
| Low Increasing | $\mu = 3$ | 0.89 |
| Low Increasing | $\mu = 10$ | 0.83 |

**Table 4.** Coverage probability of the 95% confidence interval on the mean prevalence during the middle year in an *M. bovis* cluster estimated with simple linear regression, for the different tested prevalence levels (intercept *α*; see text) and for the different trapping pressures *μ*. The value of *μ* corresponds to the mean number of animals trapped in each commune.

| Intercept | TrapPress | Probability |
| --- | --- | --- |
| $\alpha = -4$ | $\mu = 0.5$ | 0.91 |
| $\alpha = -4$ | $\mu = 1$ | 0.90 |
| $\alpha = -4$ | $\mu = 3$ | 0.78 |
| $\alpha = -4$ | $\mu = 10$ | 0.58 |
| $\alpha = -3$ | $\mu = 0.5$ | 0.94 |
| $\alpha = -3$ | $\mu = 1$ | 0.90 |
| $\alpha = -3$ | $\mu = 3$ | 0.79 |
| $\alpha = -3$ | $\mu = 10$ | 0.56 |
| $\alpha = -2$ | $\mu = 0.5$ | 0.93 |
| $\alpha = -2$ | $\mu = 1$ | 0.90 |
| $\alpha = -2$ | $\mu = 3$ | 0.80 |
| $\alpha = -2$ | $\mu = 10$ | 0.64 |
| $\alpha = -1$ | $\mu = 0.5$ | 0.92 |
| $\alpha = -1$ | $\mu = 1$ | 0.90 |
| $\alpha = -1$ | $\mu = 3$ | 0.77 |
| $\alpha = -1$ | $\mu = 10$ | 0.58 |
| $\alpha = 0$ | $\mu = 0.5$ | 0.92 |
| $\alpha = 0$ | $\mu = 1$ | 0.92 |
| $\alpha = 0$ | $\mu = 3$ | 0.86 |
| $\alpha = 0$ | $\mu = 10$ | 0.66 |

Finally, the last set of simulations showed that as long as the sample of trapped animals can be considered a random sample from the population, the model is robust to violations of the underlying hypotheses (Fig 5 and Tab 5). However, when the animals are preferentially trapped in places where the prevalence is high, the mean prevalence is overestimated (and this bias will be greater when the spatial heterogeneity is strong), and the mean proportion of animals becoming infected in one year will also be biased (although this bias is much smaller than the bias affecting the mean prevalence, and can be ignored for moderate spatial heterogeneity). Similarly, nonrandom sampling can generate bias in the estimation of the two parameters when both the spatial structure changes with time and when the sampling is directed toward highly infected communes. Note that in our study, the sampling intensity was uncorrelated with the commune random effects in the Dordogne/Charentes (Pearson correlation coefficient between the number of trapped badgers and *u*_*i*_, *R* = -0.02) and the Bearn (*R* = 0.04) clusters, whereas the trapping effort exhibited a slight preference for the most infected communes in the Burgundy cluster (*R* = 0.35).

**Figure 5.**
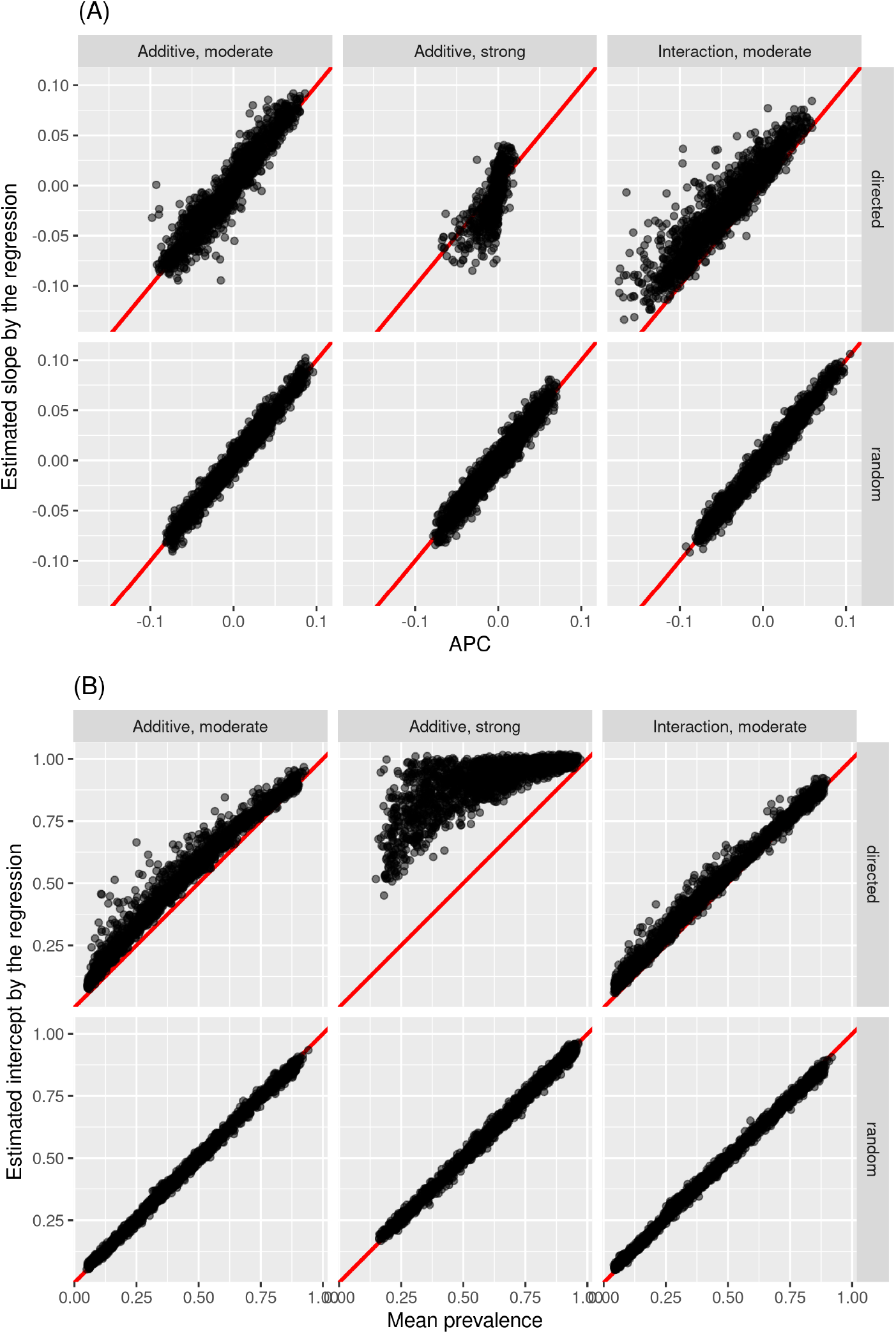
Comparison of the two statistics of interest – (A) proportion of animals becoming infected in one year, and (B) mean prevalence during the middle year – estimated using the regression indicator (see text) with the true values, estimated by simulations for the two different sampling schemes (directed = top row; random sampling = bottom row) and the different situations (additive effects of space and time on prevalence either with a moderate [*τ* = 0.73] or strong [*τ* = 0.1) spatial structure or interaction between space and time on prevalence with a moderate spatial structure [*τ* = 0.73]). Here, we pool the data simulated with the two possible values of the intercept *α* = *−*2 or *α* = 0. The straight line is the line of equation *y* = *x*.

**Table 5.**
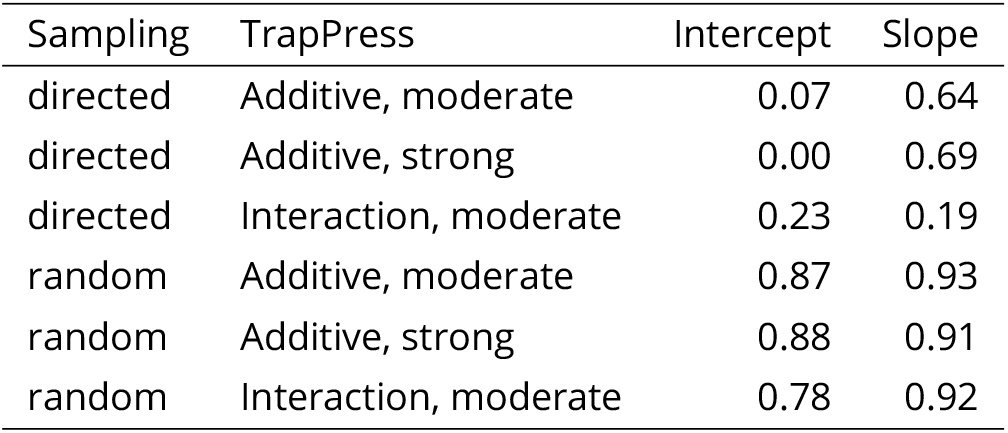
Coverage probability of the 95% confidence interval for the mean prevalence (intercept) during the middle year and the mean proportion of animals in a cluster infected in one year (slope) in an *M. bovis* cluster estimated with simple linear regression for the different sampling schemes and situations (additive effects of space and time on prevalence either with moderate [*τ* = 0.73] or strong [*τ* = 0.1) spatial structures, or interaction between space and time on prevalence with moderate spatial structure [*τ* = 0.73]).

| Sampling | TrapPress | Intercept | Slope |
| --- | --- | --- | --- |
| directed | Additive, moderate | 0.07 | 0.64 |
| directed | Additive, strong | 0.00 | 0.69 |
| directed | Interaction, moderate | 0.23 | 0.19 |
| random | Additive, moderate | 0.87 | 0.93 |
| random | Additive, strong | 0.88 | 0.91 |
| random | Interaction, moderate | 0.78 | 0.92 |

## Discussion

We developed a complex Bayesian model to describe how the infection status of badgers changed in space and time in three *M. bovis* clusters in France, accounting for the resolution of the data (commune scale), the spatial structure of the infection, the imperfect and variable sensitivity of the diagnostic tests, and the possible correlation of the infection status of badgers within the same commune. This model allowed us to estimate both the mean prevalence level and the mean proportion of badgers becoming infected in one year. We also developed an alternative, much simpler model of the infection process, based on classical linear regression, which also allowed us to easily estimate these two quantities in highly infected communes only. Simulations of the complex model showed that the two simpler indicators were good approximations of the true quantities, and could easily be used by stakeholders to estimate the key parameters of the infection process in the most infected communes.

Basically, if the tests used to diagnose *M. bovis* were characterized by a sensitivity of 100%, our regression approach would be equivalent to a simple linear regression of the *M. bovis* infection status of each animal coded as a binary variable as a function of the year (the form of the response variable *B*_*i*_/*s*_*i*_ in Equation (5) is just a way to account for the imperfect sensitivity of the tests). The suggestion to use a classical linear regression to model what is basically a binary variable can seem surprising, given that such variables are usually modeled with logistic regressions. We preferred to fit a classical linear regression since its coefficients (intercept and slope of the year) are directly interpretable as the mean prevalence level and proportion of animals becoming infected in one year respectively. Using classical linear regression to predict a binary variable leads to the violation of several hypotheses underlying this method. However, this violation is not a problem when the aim is to estimate the regression parameters, as long as we do not want to use the regression model to predict the infection status of each animal. Thus, as long as we are interested only in the slope and intercept of the regression, it does not matter that the linear regression can, in theory, predict probabilities of infection greater than 1 or lower than 0. Similarly, as noted by Gelman and Hill (2006, p. 46), “*for the purpose of estimating the regression line (as compared to predicting individual data points), the assumption of normality is barely important at all*”. Finally, the violation of the homoscedasticity assumption (equal variance of the residuals for all the predicted values) is also a minor issue in this case (Gelman and Hill, 2006, p. 46). The greater interpretability of the regression coefficients and the easier application of linear regression have led several authors to recommend this method instead of logistic regression for binary variables (Gomila, 2021; Hellevik, 2009), as long as the model is not used to predict new data points. Note, however, that the departure from the normal distribution led to low coverage probabilities for the two parameters (and especially the mean prevalence level at mid-period) when the sample size was large. Indeed, under these conditions, the departure from normality has a stronger effect on the estimation of the precision of the parameters. However, as long as the mean sample size in a commune is not too large (e.g., less than 3 animals per commune and per year), the coverage probability of the 95% confidence intervals derived from the linear regression for these parameters is close to the nominal level and can provide a rough first approximation of the uncertainty of the target quantities.

The correlation between the infection status of badgers trapped in the same commune during a given year was low (*≈* 0.03), and we showed that ignoring this correlation was not characterized by strongly biased measures of precision. Other authors have found that different badgers of the same sett have a greater chance of being infected (e.g. Delahay et al., 2000; Weber, Bearhop, et al., 2013). However, our spatial resolution was much coarser than that used in the studies of these authors: we worked at the commune scale (median area of 12 km^2^), whereas the badger home range rarely exceeds 4 km^2^ and is often much smaller (Elmeros et al., 2005; Payne, 2014). The traps set up in a commune often allow the capture of badgers from different social groups, thereby limiting the resulting correlation between infection status. Moreover, the local environmental context may be highly variable around different traps within a given commune (e.g., some places can be very close to an infected farm whereas others can be much further), which also limits this correlation. In addition, on a larger scale, in the complex multihost system encountered in France, the source of *M. bovis* infection for badgers might vary and may also come from other wild hosts, such as wild boar (whose movements may exceed the commune scale). If traps are set where interspecies transmission may occur, the correlation may be limited at a commune scale.

Our complex model identified a very marked spatial structure of the infection in the three studied *M. bovis* clusters, and both our complex model and the simpler regression approach assumed that this structure was stable in time (i.e., the areas with the highest prevalence remained the same every year; even if the mean prevalence increased or decreased in time, it changed in the same way everywhere). In statistical terms, we assumed that time and space had additive effects on the prevalence. If the spatial distribution of the infection had changed over time, which can occur for certain diseases (e.g. with some clusters becoming larger with time; see Wobeser, 1994, p. 29), this assumption would be violated. Simulations showed that a mild violation of this assumption does not impede its ability to assess the average situation in a cluster (mean prevalence and mean trend in prevalence), provided that the sample of trapped badgers can be considered entirely random, a condition we show to be approximately valid in our study (i.e., weak correlation between the sampling pressure and the prevalence of *M. bovis* infection). Moreover, this assumption of additivity is reasonable for *M. bovis* infection, as demonstrated by both a preliminary exploratory analysis of our dataset and by the epidemiological properties of this infection. The preliminary fit of a simplistic generalized additive model to predict the infection status of trapped badgers as a function of space and time showed that space-time interactions could be ignored in all clusters and that the spatial distribution of the infection in badgers was stable over time during our study period (see Appendix E for more details).

This stable spatial structure of the infection can be explained by the infection dynamics of *M. bovis* in relation to the structure of the multihost system. Indeed, infection of the badger population may result from two different dynamics: within-species transmission related to the social structure of the badger population, and between-species transmission caused by contact with infected animals of other species – in our context, mainly cattle and wild boar. The relative importance of these two dynamics varies according to the context. For instance, in Burgundy, in a recent study, we found that the spatial structure of the infected badger population was strongly related to the spatial structure of the pastures of infected cattle (Bouchez-Zacria, Payne, et al., 2023), suggesting that a between-species transmission dynamic, still active 20 years after infection, was detected in both the cattle and badger populations. In any case, within- and between-species infection dynamics logically lead to a strong and stable spatial structure of badger infection because of (i) the strong social structure of the badger population associated with a small number of dispersing animals that usually move between adjacent groups (Rogers et al., 1998); (ii) the strong spatial structure of the main external source of infection, i.e., the cattle population, which has been relatively stable over the years; and (iii) the *M. bovis* transmission mode, which involves direct or indirect contact between animals as well as an infection resulting frequently in a chronic disease (with animals being infectious for a long time). Thus, these elements suggest that the spatial diffusion of *M. bovis* infection is rather slow so it is reasonable to assume that the spatial structure of the infection in a cluster is stable over a period of a few years (e.g., 5 to 10 years). The two proposed indicators can therefore be used at this time scale to monitor changes in the infection pattern. In particular, a few informal tests of the indicators seem to indicate that a 5-year scale is an interesting scale for assessing the effect of management measures implemented to control *M. bovis* infection. When the study period covers more than 10 years, a sliding window in time can be used to fit the linear regression.

The Bayesian model accounted for the spatial structure of *M. bovis* infections in each cluster. In contrast, the regression model did not consider this spatial structure. Therefore, we recommend focusing only on highly infected communes when applying the regression model, assuming that the remaining spatial variability within this subset of communes is negligible. Note that our simulations showed that even in the presence of a substantial remaining spatial structure, there was no detectable bias in the estimation of the two focus parameters (mean proportion of the population becoming infected in one year and mean prevalence during the middle year), provided that the sample of trapped badgers could be considered completely random. When sampling is directed toward communes with the highest infection prevalence, a substantial remaining spatial structure within these highly infected communes will result in the preferential sampling of infected animals. Neglecting the spatial structure of the infection in the regression model then leads to an overestimation of the mean prevalence during the middle year and a biased estimation of the proportion of the population becoming infected in one year. Therefore, monitoring programs intending to use our regression approach should pay attention to maintaining uniform trapping pressure across a clusters’ entire area. In our study, the correlation between the level of infection in a commune and the sampling effort remained low, suggesting a very limited bias in our estimation.

We assumed equal trappability between infected and noninfected badgers. However, previous studies have shown that the trappability of badgers may be influenced by factors such as weather, season or age class (Byrne et al., 2012; Martin et al., 2017). Therefore, trappability might also vary based on other individual characteristics, and particularly the infection status of the animal, although we did not find any study supporting this hypothesis. In addition, other factors related to the infection status of badgers may indirectly affect their trappability. Thus, several studies suggest that infection can lead to behavioral changes in badgers, making them more solitary and mobile, with larger home ranges (Cheeseman and Mallinson, 1981; Garnett et al., 2005; Weber, Carter, et al., 2013). In particular, greater mobility of infected animals was observed in the three clusters in our study, leading to an increased risk of being killed by cars; the proportion of infected badgers is greater in animals killed by cars collected on the side of roads than in trapped badgers (unpublished results). This greater mobility of infected badgers may increase their exposure to traps. However, even if there was a lingering bias in the prevalence estimation, there is no indication that this bias varied among the three clusters or between years. Therefore, it is reasonable to assume that the situations can be compared consistently across clusters or between years.

During our study period, we observed different tendencies in the 3 main *M. bovis* clusters in France. In Burgundy, there was an annual decrease in the proportion of infected badgers between 2013 and 2019, and the mean prevalence in 2016 was estimated to be 0.08 (0.065-0.097) with the model whereas in the 2 other *M. bovis* clusters the tendency was either an annual increase in the proportion of infected badgers (Dordogne/Charente) or a stabilization (Bearn) with a slightly higher mean prevalence than in Burgundy: 0.126 (0.109-0.143) and 0.112 (0.092-0.134), respectively. The observations in the captured badger population are in line with the bTB situation in the bovine population. Indeed, in Burgundy, the incidence on cattle farms decreased during the same period, which was not the case for the 2 other clusters (Delavenne, Desvaux, et al., 2021). Burgundy strengthened bTB control measures earlier than did the other regions, especially in terms of early detection of infected cattle farms and in badger culling pressure, at least for some years. This is most likely the main reason for these differences, even if differences in the badger population and multihosts structures may also have played a role. Southwest of France (covering the 2 clusters with the highest proportion of infected badgers), now has the highest number of *M. bovis* cases (80% of cattle bTB cases and 94% of wildlife cases – all species included– in 2018; see Delavenne, Desvaux, et al., 2021), and additional years of effort are needed to see an improvement in epidemiological indicators.

Having a follow-up of such indicators is therefore crucial for assessing the efficiency of the measures applied. In Sylvatub, it will now be easier to reevaluate the developed indicators regularly in at-risk areas. We demonstrated that our indicators need to be calculated for the most infected communes. In our study, the complex Bayesian model that we used allowed us to identify highly infected communes (i.e., those with a random effect greater than the average); thus, these communes can be used in later monitoring for the calculation of the indicators.

If the present indicators are to be used in other situations (e.g., in newly discovered clusters or in other countries), there are several options for identifying those highly infected places. One possibility would be to fit the complex model once, a few years after the time of discovery of the cluster, to identify those communes. However, other approaches could also be used. Thus, given the reasonable additivity of space and time effects on the infection at a time scale of a few years, one could try to describe the spatial distribution of the infection risk using data collected over a short period by ignoring the time dimension. For example, the nonparametric approach of Kelsall and Diggle (1995), which estimates the spatial distribution of risk by calculating the ratio of two probability densities of positive and negative tests in space, could be used to identify more infected places.

We developed this regression approach, focusing on the badger populations in infected areas in France; however, in theory, this approach could be used more generally for any infection characterized by the additivity of space and time effects on the prevalence. Thus, the preliminary results indicate that this regression approach could also be used for wild boar in the three main French *M. bovis* clusters. In this case, the same Bayesian model provides a good description of the infection (although the spatial structure is much less clear, C. Calenge pers. com.), which suggests that the linear regression indicators proposed for the badgers could also be used for wild boar monitoring.

## Acknowledgements

We warmly thank all the stakeholders of Sylvatub, including the French Reference Laboratory for Tuberculosis (Anses, Maisons-Alfort, France), local state veterinary services, departmental hunters’ federations, the local services of the French Agency for Biodiversity, departmental laboratories for analysis, local animal health defense associations, pest control officers, and trappers and their associations. We are also indebted to Julie Rivière, the national coordinator of Sylvatub from 2011 to 2013, and the members of the National Steering Commission and Coordination Subcommittee. We extend our gratitude to Rowland Kao and an anonymous referee for their constructive comments, which significantly contributed to the enhancement of our manuscript. Special thanks to AJE for their meticulous editing, ensuring the manuscript’s linguistic excellence.

## Data, script, code, and supplementary information availability

All the data, script, codes and supplementary information have been packaged in an R package named *badgertub*, available on Github at https://github.com/ClementCalenge/badgertub. We have also stored this package on Zenodo, where it has been given the following Digital Object Identifier: https://dx.doi.org/10.5281/zenodo.10400483. The raw dataset used in this paper has also been stored as a text file on Zenodo and is available at the following URL: https://dx.doi.org/10.5281/zenodo.8010664

## References

Bouchez-Zacria M, A Courcoul, and B Durand (2018). The Distribution of Bovine Tuberculosis in Cattle Farms Is Linked to Cattle Trade and Badger-Mediated Contact Networks in South-Western France, 2007–2015. Frontiers in Veterinary Science 5. 10.3389/fvets.2018.00173.

Bouchez-Zacria M,A Payne, S Girard, C Richomme, ML Boschiroli, M Marsot, B Durand, and S Desvaux (2023). Spatial association of Mycobacterium bovis infection in cattle and badgers at the pastures interface in an endemic area in France. Under review.

Byrne AW J O’Keeffe, S Green, DP Sleeman, LAL Corner, E Gormley, D Murphy, SW Martin, and J Davenport (Dec. 2012). Population Estimation and Trappability of the European Badger (Meles meles): Implications for Tuberculosis Management. PLoS ONE 7, e50807. ISSN: 1932-6203. 10.1371/journal.pone.0050807.

Cheeseman CL and PJ Mallinson (June 1981). Behaviour of badgers (Meles meles) infected with bovine tuberculosis. Journal of Zoology 194, 284–289. ISSN: 1469-7998. 10.1111/j.1469-7998.1981.tb05780.x.

Danielsen F, MM Mendoza, P Alviola, DS Balete, M Enghoff, MK Poulsen, and AE Jensen (2003). Biodiversity monitoring in developing countries: what are we trying to achieve? Oryx 37, 407–409. 10.1017/S0030605303000735.

Delahay R, S Langton, G Smith, R Clifton-Hadley, and C Cheeseman (2000). The spatio-temporal distribution of Mycobacterium bovis (bovine tuberculosis) infection in a high-density badger population. Journal of Animal Ecology 69, 428–441. 10.1046/j.1365-2656.2000.00406.x.

Delavenne C, F Pandolfi, S Girard, E Réveillaud, P Jabert, M Boschiroli, L Dommergues, F Garapin, N Keck, F Martin, M Moussu, S Philizot, J Rivière, I Tourette, D Calavas, C Dupuy, B Dufour, and F Chevalier (2019). Tuberculose bovine : bilan et évolution de la situation épidémiologique entre 2015 et 2017 en France métropolitaine. Bulletin épidémiologique 91, 1–22.

Delavenne C, S Desvaux, ML Boschiroli, S Carles, P Chaigneau, B Dufour, B Durand, K Gache, F Garapin, S Girard, P Jabert, N Keck, É Réveillaud, J Rivière, C Dupuy, and F Chevalier (2021). Surveillance de la tuberculose due à mycobacterium bovis en France métropolitaine en 2018 : résultats et indicateurs de fonctionnement. Bulletin Épidémiologique 94, 1–9.

Dohoo I, W Martin, and H Stryhn (2003). Veterinary epidemiologic research. University of Prince Edward Island, p. 706.

Elmeros M, AB Madsen, and A Prang (2005). Home range of the badger (Meles meles) in a heterogeneous landscape in Denmark. Lutra 48, 35.

Garnett B, R Delahay, and T Roper (Oct. 2005). Ranging behaviour of European badgers (Meles meles) in relation to bovine tuberculosis (Mycobacterium bovis) infection. Applied Animal Behaviour Science 94, 331–340. ISSN: 0168-1591. 10.1016/j.applanim.2005.02.013.

Gelman A and X Meng (1996). Model checking and model improvement. In: Markov chain Monte Carlo in practice. Ed. by Gilks W and Richardson S. Chapman & Hall/CRC. Chap. 11, pp. 189–201.

Gelman A and D Rubin (1992). Inference from iterative simulation using multiple sequences. Statistical Science 7, 457–472. 10.1214/ss/1177011136.

Gelman A and J Hill (2006). Data analysis using regression and multilevel/hierarchical models. Cambridge University Press.

Gelman A and I Pardoe (2007). 2. Average predictive comparisons for models with nonlinearity, interactions, and variance components. Sociological Methodology 37, 23–51. 10.1111/j.1467-9531.2007.00181.x.

Gomila R (2021). Logistic or Linear? Estimating Causal Effects of Experimental Treatments on Binary Outcomes Using Regression Analysis. Journal of Experimental Psychology 150, 700–709. 10.1037/xge0000920.

Hellevik O (2009). Linear versus logistic regression when the dependent variable is a dichotomy. Quality & Quantity 43, 59–74. 10.1007/s11135-007-9077-3.

Hisakado M, K Kitsukawa, and S Mori (2006). Correlated binomial models and correlation structures. Journal of Physics A: Mathematical and General 39, 15365. 10.1088/0305-4470/39/50/005.

Hosmer D and S Lemeshow (2000). Applied logistic regression. Second Edition. John Wiley & Sons.

Jacquier M, JM Vandel, F Léger, J Duhayer, S Pardonnet, L Say, S Devillard, and S Ruette (2021). Breaking down population density into different components to better understand its spatial variation. BMC Ecology and Evolution 21. 10.1186/s12862-021-01809-6.

Kelsall JE and PJ Diggle (1995). Non-parametric estimation of spatial variation in relative risk. Statistics in medicine 14, 2335–2342. 10.1002/sim.4780142106.

King G and L Zeng (2002). Estimating risk and rate levels, ratios and differences in case-control studies. Statistics in medicine 21, 1409–1427. 10.1002/sim.1023.

Marsot M, M Béral, A Scoizec, Y Mathevon, B Durand, and A Courcoul (2016). Herd-level risk factors for bovine tuberculosis in French cattle herds. Preventive Veterinary Medicine 131, 31–40. 10.1016/j.prevetmed.2016.07.006.

Martin LER, AW Byrne, J O’Keeffe, MA Miller, and FJ Olea-Popelka (Feb. 2017). Weather influences trapping success for tuberculosis management in European badgers (Meles meles). European Journal of Wildlife Research 63. ISSN: 1439-0574. 10.1007/s10344-017-1089-2.

Palisson A, A Courcoul, and B Durand (Mar. 2016). Role of Cattle Movements in Bovine Tuberculosis Spread in France between 2005 and 2014. PLOS ONE 11, 1–19. 10.1371/journal.pone.0152578.

Payne A (2014). Rôle de la faune sauvage dans le système multi-hôtes de Mycobacterium bovis et risque de transmission entre faune sauvage et bovins: étude expérimentale en Côte d’Or. PhD thesis. Université Claude Bernard-Lyon I.

Plummer M, N Best, K Cowles, and K Vines (2006). CODA: Convergence Diagnosis and Output Analysis for MCMC. R News 6, 7–11.

Pocock MJ, SE Newson, IG Henderson, J Peyton, WJ Sutherland, DG Noble, SG Ball, BC Beckmann, J Biggs, and T Brereton (2015). Developing and enhancing biodiversity monitoring programmes: a collaborative assessment of priorities. Journal of Applied Ecology 52, 686–695. 10.1111/1365-2664.12423.

R Core Team (2023). R: A Language and Environment for Statistical Computing. R Foundation for Statistical Computing. Vienna, Austria.

Réveillaud E, S Desvaux, ML Boschiroli, J Hars, É Faure, A Fediaevsky, L Cavalerie, F Chevalier, P Jabert, S Poliak, I Tourette, P Hendrikx, and C Richomme (2018). Infection of Wildlife by Mycobacterium bovis in France Assessment Through a National Surveillance System, Sylvatub. Frontiers in Veterinary Science 5, 262. 10.3389/fvets.2018.00262.

Riviere J, Y Le Strat, B Dufour, and P Hendrikx (2015). Sensitivity of bovine tuberculosis surveillance in wildlife in France: a scenario tree approach. PLoS One 10, e0141884. 10.1371/journal.pone.0141884.

Rivière J, J Hars, C Richomme, A Fediaevsky, D Calavas, E Faure, and P Hendrikx (2012). La surveillance de la faune sauvage : de la théorie à la pratique avec l’exemple du réseau Sylvatub. Épidémiologie et Santé Animale 61, 5–16.

Rogers L, R Delahay, C Cheeseman, S Langton, G Smith, and R Clifton-Hadley (1998). Movement of badgers (Meles meles) in a high–density population: individual, population and disease effects. Proceedings of the Royal Society of London B: Biological Sciences 265, 1269–1276. 10.1098/rspb.1998.0429.

Roper TJ (2010). Badger. HarperCollins UK.

Rue H and L Held (2005). Gaussian Markov Random Fields. Theory and Applications. Chapman & Hall/CRC.

Valpine P de, D Turek, CJ Paciorek, C Anderson-Bergman, DT Lang, and R Bodik (2017). Programming with models: writing statistical algorithms for general model structures with NIMBLE. Journal of Computational and Graphical Statistics 26, 403–413. 10.1080/10618600.2016.1172487.

Weber N, S Bearhop, SR Dall, RJ Delahay, RA McDonald, and SP Carter (2013). Denning behaviour of the European badger (Meles meles) correlates with bovine tuberculosis infection status. Behavioral Ecology and Sociobiology 67, 471–479. 10.1007/s00265-012-1467-4.

Weber N, SP Carter, SR Dall, RJ Delahay, JL McDonald, S Bearhop, and RA McDonald (Oct. 2013). Badger social networks correlate with tuberculosis infection. Current Biology 23, R915–R916. ISSN: 0960-9822. 10.1016/j.cub.2013.09.011.

Wickham H (2016). ggplot2: Elegant Graphics for Data Analysis. Springer-Verlag New York.

Wickham H and G Grolemund (2017).R for Data Science: Import, Tidy, Transform, Visualize, and Model Data. 1st ed. O’Reilly Media.

Wobeser GA (1994). Investigation and management of disease in wild animals. Springer Science & Business Media.

