## Supplementary material for "Assessing the dynamics of *Mycobacterium bovis* infection in three French badger populations"

### Introduction

This vignette is the supplementary material of the article of Calenge et al. (in prep.). The aim of this paper is to model the *Mycobacterium bovis* infection in three French badger populations between 2013 and 2019, and to propose two new indicators for the monitoring of the prevalence mean level and trend in each population. A companion package named `badgertub` contains the data and functions used for this paper, and is required to reproduce the calculations in this document. The present document is also available as a vignette of this package. To install this package, first install the package `devtools` and use the function `install_github` to install `badgertub`:

```
## If devtools is not yet installed, type
install.packages("devtools")

## Install the package badgertub
devtools::install_github("ClementCalenge/badgertub", ref="main")
```

*Remark*: on Windows, it is required to also install the Rtools (`https://cran.r-project.org/bin/windows/Rtools/`) on your computer to have a working `devtools` package (see `https://www.r-project.org/nosvn/pandoc/devtools.html`).

Throughout this vignette, we suppose that the reader is familiar with the model and simulations developed in the main paper. More precisely:

- In appendix A, we give additional information regarding the formal expression of the beta-binomial distribution used in the model.
- In appendix B, we give additional information regarding the average predictive comparison calculated to assess the effect of the year in our model of the prevalence.
- In appendix C, we describe how we derived our two indicators of the prevalence level and trend.
- In appendix D, we give the R code used to fit the model and simulate virtual situations to test the indicators, as well as additional checks of the fitted models (goodness of fit, MCMC convergence).
- In appendix E, we describe a preliminary modelling approach relying on generalized additive models that we used as a first exploratory approach to confirm the additivity of space and time effects.

### Appendix A The beta-binomial distribution

Our model suppose that the number $y_{it}$ of animals actually infected among the $N_{it}$ trapped and tested animals in commune $i$ during year $t$ can be described by a beta-binomial distribution:

$$y_{it} \sim \text{Beta-Binomial}(N_{it}, p_{it}, \rho)$$

Where $p_{it}$ is the true mean prevalence in commune $i$ during year $t$, and $\rho$ is the correlation between the infection status of two badgers randomly drawn in our dataset (both $p_{it}$ and $\rho$ are estimated by the model fit).

In this appendix, we describe the parameterization of the beta distribution as a function of the prevalence and correlation between animals.

The beta-binomial distribution is a commonly used distribution to account for a correlation between Bernoulli variables (Hisakado et al., 2006; Martin et al., 2011). A variable $y$ follows a beta-binomial distribution if $y$ follows a binomial distribution characterized by a size $N$ and a probability $p$ randomly distributed according to a beta distribution with parameters $\alpha$ and $\beta$. The probability of a beta-binomial variable $y$ is therefore:

$$P(y|N, \alpha, \beta) = \left( \begin{array}{c} N \\ y \end{array} \right) \frac{\mathrm{B}(y + \alpha, N - y + \beta)}{\mathrm{B}(\alpha, \beta)}$$

With $\left( \begin{array}{c} N \\ y \end{array} \right)$ the binomial coefficient, and the beta function $\mathrm{B}(u, v)$ is calculated by:

$$B(u, v) = \frac{\Gamma(u)\Gamma(v)}{\Gamma(u + v)}$$

and $\Gamma(x)$ is the gamma function (extension of the factorial function).

Note that this distribution is characterized by an expectation equal to:

$$\mu = Np'$$

with $p'$ the mean probability, calculated by:

$$p' = \frac{\alpha}{\alpha + \beta}$$

In our model, this parameter $p'$ is the mean prevalence $p_{it}$ modelled as a function of the year and commune in our approach.

Hisakado et al. (2006) show that the beta-binomial distribution can be used to model the correlation between the infection status of two animals randomly drawn in the population. Indeed, if $y$ follows a beta-binomial distribution, then the infection status of two randomly drawn animals is correlated, with a correlation coefficient equal to:

$$\rho = \frac{1}{\alpha + \beta + 1}$$

In other words, the beta-binomial distribution can be considered as a particular distribution corresponding to the sum of $N$ correlated Bernoulli variables.

We can parameterize the beta-distribution to make this point of view clearer. Thus, we represent this distribution as a distribution parameterized by $N, p', \rho$. Note that we can easily transform back this distribution to find the parameters $\alpha$ and $\beta$ of the beta distribution from the parameters $p', \rho$:

$$\begin{aligned} \alpha &= p' \times \frac{(1 - \rho)}{\rho} \\ \beta &= (p' \times \rho - p' - \rho + 1)/\rho \end{aligned}$$

We can fit the beta-binomial distribution either with the parameters $p', \rho$, or with the parameters $\alpha, \beta$.

### Appendix B The average predictive comparison (APC)

In this appendix, we give additional details on the calculation of the APC. Recall that the APC of a variable $u$ in a given model corresponds to the **predicted mean difference of the response variable when the predictive variable $u$ increases by one unit, while other predictive variables are being held constant**. For example, when the model is a simple or multiple linear regression without interactions, the APC of a predictive variable is simply the slope of the variable. However, when the model is more complex (e.g. when the variable $u$ interacts with other variables, or when the model is nonlinear such as a logistic regression), it is more difficult to estimate this "average slope".

For example, the coefficient of a variable $u$ in a logistic regression describes how the *logit* of the response variable itself varies in average when $u$ increases by one unit. However, this coefficient does not bring any direct information about how the response variable varies in average for a unit increase in $u$.

In our study, $u$ is the variable year, and our aim is to calculate the expected difference of the probability of infection when one year passes (i.e. the proportion of the population becoming infected in one year), which is precisely what the APC of the year estimates in the logistic model predicting the prevalence of *M. bovis* infection.

We therefore use this approach. We now describe precisely how to estimate the APC for any type of model. Let $y$ be the response variable (in our study, the infected/non-infected status of animals). Following Gelman and Pardoe (2007), we define:

- $u$ the predictive variable for which the APC is to be estimated (in our study, the year).

- $v$ all the other predictive variables in the model (in our study, the commune where the animal was trapped).

Let $\theta$ be the vector of parameters of the model. As noted in the paper, the *predictive comparison* (PC) associated to a variable $u$ in a model is defined by:

$$\delta_u(u^{(1)} \to u^{(2)}, v, \theta) = \frac{\mathrm{E}(y|u^{(2)}, v, \theta) - \mathrm{E}(y|u^{(1)}, v, \theta)}{u^{(2)} - u^{(1)}}$$

This PC describes how the response variable varies when the predictive variable of interest varies from $u^{(1)}$ to $u^{(2)}$, the value of other predictive variables $v$ and the model parameters $\theta$ being held constant. When we work with a model with a known structure, and when the parameters $\theta$ are known, and when the variables $v$ have a known value, we can easily calculate the expectation of the response variable $\mathrm{E}(y|u, v, \theta)$ from the model. Therefore, for a vector of parameters $\theta$ and given values $v$, it is easy to calculate the PC of the variable in the model.

The average PC – or APC – $\Delta_u$ corresponds to the mean of $\delta_u$, calculated over the statistical distribution of predictive variables used to fit the model and over the posterior distribution of the parameters. Formally:

$$\Delta_u = \frac{\int\int_{u^{(1)}<u^{(2)}} du^{(1)} du^{(2)} \int dv \int d\theta \{E(y|u^{(2)}, v, \theta) - E(y|u^{(1)}, v, \theta)\} p(u^{(1)}|v) p(u^{(2)}|v) p(v) p(\theta)}{\int\int_{u^{(1)}<u^{(2)}} du^{(1)} du^{(2)} \int dv \int d\theta \{u^{(2)} - u^{(1)}\} p(u^{(1)}|v) p(u^{(2)}|v) p(v) p(\theta)} \quad (1)$$

We therefore consider, in our dataset, all the pairs of transitions from $(u^{(1)}, v^{(1)})$ to $(u^{(2)}, v^{(2)})$, such that $v^{(1)} = v^{(2)}$. In other words, we consider all the pairs of statistical units of the dataset (in our case, the badgers) for which the predictive variables $v$ (the communes) included in the model other than the variable of interest $u$ (the year) are held constant.

This is theory. Now, Gelman and Pardoe (2007) describe how to calculate $\Delta_u$ in practice: the MCMC samples provide a statistical distribution for the parameters, i.e. $p(\theta)$ in equation (1), and the dataset is used to estimate the distribution of the predictive variables $p(v)$ and $p(u|v)$ in this equation.

Thus, one draws $S$ random vectors of parameters $\theta^{(s)}$ from the posterior distribution of the distribution (with $S$ large, e.g. equal to 1000). If $n$ is the number of individuals in the dataset, then one can estimate the APC with:

$$\hat{\Delta}_u = \frac{\sum_{i=1}^{n}\sum_{j=1}^{n}\sum_{s=1}^{S} w_{ij}\{E(y|u_j, v_i, \theta^s) - E(y|u_i, v_i, \theta^s)\}\text{sign}(u_j - u_i)}{\sum_{i=1}^{n}\sum_{j=1}^{n}\sum_{s=1}^{S} w_{ij}\{u_j - u_i\}\text{sign}(u_j - u_i)} \tag{2}$$

In other words, this approach consists in averaging, over all possible pairs of statistical units in the dataset $i$ and $j$, the difference between the values of the response variable predicted by the model $E(y|u_j, v_i, \theta^s) - E(y|u_i, v_i, \theta^s)$ for the values of the predictive variable $u$ for these two units, while constraining these expectations to be conditional on the values of the other predictive variables $v$ observed for one of the two units, $i$.

The difficulty in the development of an estimator for the APC is that equation (1) relies on both $p(u^{(1)}|v)$ and $p(u^{(2)}|v)$ – the probability to observe variable $u = u^{(1)}$ and $u = u^{(2)}$ respectively given a value of $v$. But using the dataset, for a given value of $v = v_i$ we have a value $u = u_i$, but we do not have $(u_i, v_j)$. In other words, a given badger trapped during a given year $(u_i)$ is trapped in a given commune $v_i$, but this given badger is not trapped the same year in another commune $v_j$. In other words, if we set $v = v_i, u^{(1)} = u_i$ and $u^{(2)} = u_j$, the data can be used to approximate $p(u^{(1)}|v) = p(u_i|v_i)$ but not $p(u^{(2)}|v) = p(u_j|v)$ (see Gelman and Pardoe, 2007, for further details)

This is the reason why the estimator includes weights $w_{ij}$, which describe the proximity between $v_i$ and $v_j$. The difference between predicted values based on the pairs $(i, j)$ has larger weight in the calculation of $\hat{\Delta}_u$ when the other predictive variables $v$ are similar. The idea is to approximate $p(u^{(2)}|v)$ by giving more weight to pairs of individuals with values of $v$ that are close to one another (so that $p(u_j|v_i) \approx p(u_i|v_i)$).

In our study, we defined these proximity weights by:

$$w_{ij} = \frac{1}{1 + d_{ij}}$$

with $d_{ij}$ the Euclidean distance between the centroid of the communes where the animals $i$ and $j$ were trapped. This weighting ensure a larger weight for closer communes. We used this equation to estimate the APC of the year for our models.

### Appendix C Rationale underlying the development of the indicators of prevalence trend and mean levels

#### C.1 On the use of linear regression to estimate the mean proportion of animals becoming infected in one year

In this appendix, we give more details on the rationale underlying the use of the simple linear regresssion to estimate the mean prevalence level and the mean prevalence trend during a given period in a given cluster.

Our study of the infection in three clusters makes clear the following results: (i) the space and time effects on the prevalence are additive (i.e. the highly infected places in a given cluster are the same all along the study period, see discussion of the paper and appendix E); (ii) the correlation between actual infection status of two animals trapped in the same commune is very low and can be ignored as a first approximation; (iii) the effect of the year on the prevalence is linear (see also appendix E); (iv) there is a large spatial variability of the prevalence during a given year.

Let us ignore for the moment the variability with time of the sensitivity of the tests used to detect *M. bovis* in badgers. Let us focus only on the badgers trapped in the highly infected communes (i.e. the communes for which the spatial effect estimated with the Bayesian model is greater than the mean spatial effect). As noted in the paper, by focusing only the highly infected communes, we hope that the remaining spatial variability in prevalence is negligible (which is confirmed by the simulations). Then, in this situation, the changes with time of the epidemiological situation can be summarized by a logistic regression of the infection status of trapped badger as a function of the year. And the slope of the year in such a model completely summarizes how the infection varies with time in the cluster.

However, as noted in the paper (and in appendix B), the slope of a variable in a logistic regression can be difficult to understand as such by nonspecialists. For this reason, several authors have suggested to replace logistic regression by classical Gaussian linear regression when the only goal is to estimate an effect of predictive variables that must be easy to understand (Hellevik, 2009; Gomila, 2020). Such an approach is extremely simple to implement and its coefficients are much more interpretable. For example, a slope for the year equal to 0.02 in a linear regression predicting the infection status (1 = infected / 0 = non-infected) as a function of the year indicates that the proportion of infected animals increases by 2% per year.

#### C.2 Accounting for the imperfect sensitivity of the tests

However, we need to account for the imperfect sensitivity of the tests in the linear regression. Consider $N$ trapped badgers. Let $t_i$ be the year when the animal $i$ has been trapped, $V_i$ the *true* infection status of animal $i$, and $B_i$ the infection status detected by the test used on the individual $i$. As noted in the previous section, if we knew the true infection status of all animals, we would fit the following linear regression:

$$V_i = a + b \times t_i + \epsilon_i \tag{3}$$

with $\epsilon_i$ a residual, and $a, b$ respectively the intercept and the slope of the regression line. The coefficient $b$ would therefore be an estimator of the mean proportion of the population becoming infected in one year in the highly infected communes of the cluster.

However, the variable $V_i$ is a hidden variable: we do not know the true infection status of animal $i$ – only the detected status $B_i$.

Actually, the variable $B_i$ is a Bernoulli variable:

$$B_i = \mathcal{B}(V_i, s_i)$$

and its expectation is equal to:

$$E(B_i) = V_i s_i$$

We can therefore simply replace $V_i$ by $B_i/s_i$ in equation (3). Indeed, $E(B_i/s_i) = V_i$, which suggests that the use of this variable as a response variable in a linear regression would allow to avoid a bias caused by the test sensitivity. Our linear regression therefore becomes:

$$B_i/s_i = a + b \times t_i + \epsilon_i \tag{4}$$

#### C.3 The mean prevalence trend

We have showed that the slope of the linear regression (4) can be used as an indicator of the prevalence trend. However, we can also use the intercept of the regression as an indicator of the prevalence level. Let us recall that in a classical linear regression, the intercept corresponds to the expected value of the response variable when the predictive variables are equal to 0. Therefore, if we modify slightly the linear regression:

$$B_i/s_i = a + b \times \tilde{t_i} + \epsilon_i \tag{5}$$

where $\tilde{t_i}$ is the centred year (i.e. $t_i - \bar{t}$, where $\bar{t}$ is the middle year of the study period), then $\tilde{t_i} = 0$ corresponds to the middle year of the study period, so that the coefficient $a$ measures the expected value of the prevalence during the middle year of the study period.

Thus, the slope and intercepts of equation (5) can be used as indicators of the prevalence trend and level respectively.

### Appendix D R code used to fit the model and for the simulations

In this appendix, we provide the R code used to fit the models describing the infection of badgers by *M. bovis* in our paper, as well as the simulations carried out to test the proposed indicators.

#### D.1 Model fit

##### D.1.1 The data

We first show how we fitted the models in our study. We illustrate the approach with the Dordogne/Charentes cluster, but the reader can easily reproduce the same approach for the two other clusters, as the Sylvatub data are also provided for these clusters.

First we load the package:

```
library(badgertub)
```

And then, we load the data for this cluster:

```
## Two maps
data(dpt)
data(cfcdo)

## The Sylvatub dataset
data(dotub)

str(dotub)

## List of 2
##  $ data  :List of 7
##   ..$ adj        : int [1:2452] 30 36 39 80 92 102 47 63 71 137 ...
##   ..$ num        : int [1:444] 6 4 4 6 3 4 6 6 10 5 ...
##   ..$ weights    : num [1:2452] 1 1 1 1 1 1 1 1 1 1 ...
##   ..$ Nb         : int [1:1509] 3 2 1 7 1 1 3 1 1 3 ...
##   ..$ y          : int [1:1509] 0 0 0 0 0 0 0 0 0 0 ...
##   ..$ sensitivity: num [1:1509] 0.5 0.5 0.75 0.75 0.75 0.5 0.75 0.75 0.5 0.5 ...
##   ..$ year       : num [1:1509] -2.2 -1.2 -0.2 0.8 2.8 -2.2 -0.2 0.8 -1.2 -1.2 ...
##  $ consts:List of 4
##   ..$ N : int 1509
##   ..$ P : int 444
##   ..$ L : int 2452
##   ..$ ID: Named int [1:1509] 1 1 1 1 1 2 2 2 3 4 ...
##   .. ..- attr(*, "names")= chr [1:1509] "16004" "16004" "16004" "16004" ...
```

The dataset `dpt` is an object of class `"sf"` (from the package `sf`) containing the map of the departments (French administrative division); the dataset `cfcdo` is also an object of class `"sf"` containing the map of the communes of the Dordogne/Charentes Cluster. The interested reader can plot with the function `plot()` to see these maps.

Finally, the dataset `dotub` is a list storing the Sylvatub data collected in this cluster. More precisely, this list contains two sublists, formatted to be directly usable by the package `nimble`. The sublist named `data` contains the following elements:

- `y`: For a given commune and a given year, the number of trapped badgers that were analyzed by the Sylvatub network and were identified as positive.

- `Nb`: For a given commune and a given year, the number of trapped badgers that were analyzed by the Sylvatub network.

- `sensitivity`: For a given commune and a given year, the sensitivity of the tests used (either 0.5 for a bacterial culture, or 0.75 for a PCR test).

- `year`: The centred year (i.e. with mean=0) used to fit the model.

- `adj`: a vector of indices of the adjacent communes (neighbors) of each commune. This is a sparse representation of the full adjacency matrix. This vector is used in `dcar_normal()` from the package `nimble` (see the help page of this function).

- `num`: a vector giving the number of neighboring communes of each commune in the cluster, with length equal to the total number of communes. This vector is also used in `dcar_normal()` from the package `nimble` (see the help page of this function).

- `weights`: a vector of weights, containing the symmetric unnormalized weights associated with each pair of adjacent locations, of the same length as `adj`. This vector is also used in `dcar_normal()` from the package `nimble` (see the help page of this function).

The sublist named `consts` contains the constants used for the model fit, i.e.:

- `N`: The number of commune-years in the dataset (number of elements of `y`, `Nb`, etc., in the component `data` of this list).

- `P`: The number of communes of the cluster.

- `L`: The number of elements in `adj` (in the component `data` of this list).

- `ID`: The commune ID for each element in the the component `data` of this list (`y`, `Nb`, etc.)

**Remark**: The same data are available for the two other clusters. Thus, `cfcbo` and `botub` are respectively the map of the communes and Sylvatub dataset for the Burgundy cluster. Similarly, `cfcbe` and `betub` are respectively the map of the communes and Sylvatub dataset for the Bearn cluster. The reader can reproduce the model fit for the other two clusters by just replacing `cfcdo` and `dotub` in the code below by these other datasets if they want to reproduce the analyses for the other clusters.

##### D.1.2 Model fit

We first load the package nimble, required for the fit.

```
library(nimble)
```

We then program the model with the script language used by the package `nimble`:

```
Tubcode <- nimbleCode({
    tau ~ dgamma(1,1)
    slopeYear ~ dnorm(0, 0.01)
    intercept ~ dnorm(0, 0.01)
    phi ~ dgamma(0.1,0.1)

    si[1:P] ~ dcar_normal(adj[1:L], weights[1:L], num[1:P], tau)

    for (i in 1:N) {
        lpim[i] <- intercept + si[ID[i]] + year[i]*slopeYear
        pim[i] <- exp(lpim[i])/(1+exp(lpim[i]))

        a[i] <- pim[i] * phi
        b[i] <- phi *(1-pim[i])

        p[i] ~ dbeta(a[i], b[i])
        y[i] ~ dbinom(prob=p[i]*sensitivity[i],size=Nb[i])
    }
})
```

Then, we define starting values for the parameters of the model:

```
dotubInits <- list(tau = 0.1, slopeYear = 0.2,
                   intercept=-3, phi=20,
                   si=rnorm(dotub$consts$P),
                   p=rep(0.1, dotub$consts$N))
```

And finally, we use the function `nimbleMCMC` from the package `nimble` to fit this model, directly passing the components `data` and `consts` as arguments to this function. We sample 4 chains of 1000000 MCMC samples after a burn-in period of 3000 samples. To save some disk space, we thin the chain by recording one sample every 1000. WARNING: THIS CALCULATION TAKES A VERY LONG TIME (several hours) !!!! Note that we have included the results of this calculation as a dataset of the package, so that the reader does not need to launch this function to reproduce further calculations:

```
set.seed(777)
do.mcmc.out <- nimbleMCMC(code = Tubcode, constants = dotub$consts,
                          data = dotub$data, inits = dotubInits,
                          nchains = 4, niter = 1003000, nburnin=3000, thin=1000,
                          summary = TRUE, WAIC = TRUE,
                          monitors = c('tau',"intercept","slopeYear","phi", "si"))
```

The results of this fit are available in the dataset `do.mcmc.out` of the package:

```
data(do.mcmc.out)
```

**Remark:** Similarly, the result of the fit for the Burgundy and Bearn clusters are available in the datasets `bo.mcmc.out` and `be.mcmc.out` respectively.

We convert the results to the class `"mcmc.list"` (package `coda`) to examinate the convergence of the fit. We use the function `nimbleToCoda()` available in the package `badgertub`:

```
samdo <- (do.mcmc.out$samples)
ml <- nimbleToCoda(samdo, start=3001, end=1003000, thin=1000)
```

The reader can check visually the convergence of the chain by plotting these elements (we omit the results, to save space in this document):

```
plot(ml)
```

We can check more formally this convergence for the parameters of the model with the diagnostic of Gelman and Rubin (1992) (similarly, we omit the results, to save space in this document):

```
library(coda)
gelman.diag(ml, multivariate=FALSE)
```

We just show these diagnostics for the top parameters of the model:

```
mlb <- ml
for (i in 1:4)
    mlb[[i]] <- mlb[[i]][,1:3]
gelman.diag(mlb,multivariate=FALSE)

## Potential scale reduction factors:
##
##             Point est. Upper C.I.
## phi                  1       1.01
## slopeYear            1       1.01
## tau                  1       1.01
```

These diagnostics are all lower than 1.1 (including those of the commune effects, not shown here), as recommended by Gelman and Rubin (1992). The chain convergence is satisfying.

##### D.1.3 Goodness of fit

We used the approach of Gelman and Meng (1996) to check the goodness of fit of the model. As noted in the paper, we simulated a replication of the Sylvatub dataset for each MCMC sample. We programmed a function to perform these simulations, named `simulateInfectionModel()` (see the help page of this function). We now perform these simulations. THIS CALCULATION TAKES A VERY LONG TIME !!!! Note that we have included the results of this calculation as a dataset of the package, so that the reader does not need to launch this function to reproduce further calculations:

```
simDor <- simulateInfectionModel(samdo, dotub$consts, dotub$data)
```

We load the results of these simulations:

```
data(simDor)
```

**Remark:** Similarly, the simulations carried out for the Burgundy and Bearn clusters are available in the datasets `simBou` and `simBea` respectively.

We can now use the function `compareSimActual()` of the package to compare the statistics characterizing the dataset with the same statistics calculated on the simulated datasets. More precisely: (i) for each combination commune/year, we calculated the 90% credible interval on the number of animals detected as infected expected from the model (using the distribution of simulated datasets), and we calculated the proportion of commune/years for which the observed number fell in this theoretical interval; (ii) we then considered the total number of animals detected as infected in each commune over the 7 years of the study and similarly we calculated the proportion of communes for which the observed value fell in the

90% credible interval calculated from this distribution; (iii) we calculated the total number of animals detected as infected over all communes of the cluster for each year and calculated the proportion of years for which the observed value fell in the 90% credible interval expected under the model; (iv) we calculated the 90% credible interval expected under the model on the total number of animals detected as infected over all the communes and the years, and compared it to the observed value. We show the results of these comparison below:

```
(csa <- compareSimActual(simDor, dotub$data, dotub$consts))

## **********
## ** Goodness of fit of the model
##
## Total number of infected animals:
##  obs = 247    Expected 90% CI = [216,291]
##
## Number of infected animals per year
##  Proportion of years in the 90% CI: 0.8571429
##
## Number of infected animals per commune
##  Proportion of commune in the 90% CI: 0.9951574
##
## Number of infected animals per year-commune
##  Proportion of year-commune in the 90% CI: 0.9867462
```

All these comparisons show a correct goodness of fit of the model.

##### D.1.4 Interpretation of the model

We can now use the function `summaryModel()` of the package `badgertub` to show a table of the estimated parameters in our model:

```
summaryModel(samdo)

##           Parameter          Estimates
## 1 Slope of the year 0.18  [0.11, 0.25]
## 2               Rho 0.04  [0.02, 0.08]
## 3       1/sqrt(tau)  1.17  [0.87, 1.5]
```

This table corresponds to the first three rows of Tab. 1 of the paper. We can also use the function `showSpatialEffects()` to draw a map of the spatial effects estimated by our model:

```
showSpatialEffects(samdo,cfcdo,dpt)
```

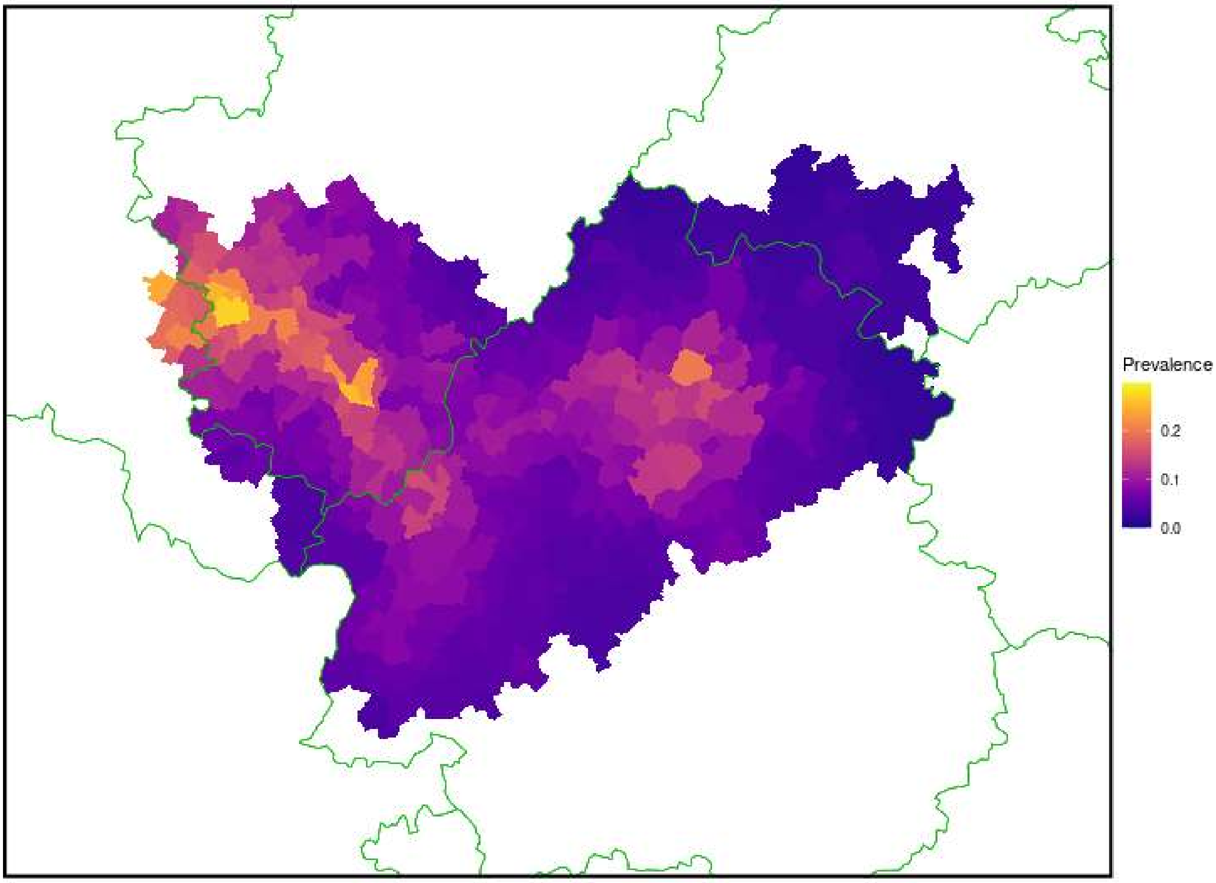

This map corresponds to Fig. 1 of the paper. We can then use the function `apc()` to estimate the average predictive comparison in our model. WARNING !!! THIS CALCULATION TAKES A VERY LONG TIME !!!! Note that we have included the results of this calculation as a dataset of the package, so that the reader does not need to launch this function to reproduce further calculations:

```
APCdo <- apc(samdo, cfcdo, dotub$consts, dotub$data)
```

We load the dataset containing the results:

```
data(APCdo)
APCdo

## *** Average predictive comparison for the year
##     Delta_u          SE
## 0.018164915 0.005386693
```

**Remark:** Similarly, the APC calculated for the Burgundy and Bearn clusters are available in the datasets `APCbo` and `APCbe` respectively.

This value corresponds to the fourth row of Tab. 1 in the paper. We can also calculate the average prevalence level in highly infected communes, using the function `averageLevel()` of the package:

```
aldo <- averageLevel(samdo,dotub$data, dotub$consts)
aldo

## *** Mean prevalence
##   MeanPrev         SE
## 0.12575172 0.01056688
```

This value corresponds to the sixth row of Tab. 1 in the paper. Finally, we can use a simple linear regression to derive the prevalence level and trends indicators. We first need to transform our dataset to a dataframe with one row per animal (thanks to the function `nimbleData2df()` of the package), and then to implement the regression approach described in the paper:

```
dfb <- nimbleData2df(dotub$data, dotub$consts, sam=samdo)
dfb$year <- dfb$year-4
dfb$response <- dfb$y/dfb$sensitivity
me <- lm(response~year, data=dfb)
summary(me)

##
## Call:
## lm(formula = response ~ year, data = dfb)
##
## Residuals:
##      Min       1Q   Median       3Q      Max
## -0.24371 -0.18618 -0.12865 -0.09988  1.92888
##
## Coefficients:
##             Estimate Std. Error t value Pr(>|t|)
## (Intercept) 0.157415   0.010071  15.630  < 2e-16
## year        0.028766   0.005198   5.534 3.51e-08
##
## Residual standard error: 0.4653 on 2160 degrees of freedom
## Multiple R-squared:  0.01398,Adjusted R-squared:  0.01352
## F-statistic: 30.62 on 1 and 2160 DF,  p-value: 3.514e-08
```

Which corresponds to the fifth and seventh row of Tab. 1 in the paper.

#### D.2 Simulations

In this section, we show the code used for the simulations carried out in the paper to demonstrate the usefulness of our indicators.

##### D.2.1 First set of simulations

The first set of simulations was carried out to assess the ability of the regression model to estimate the prevalence trend in two different situations: (i) low but increasing prevalence, and (ii) high prevalence, increasing or decreasing. We define the parameters of the simulations below (the comments in the R code explain the parameters):

```
## A list of length two (one element per simulated situation).  Each
## element is a vector containing the limits of the range within which
## the slope of the year is randomly sampled:
```

```r
##
## - Situation 1: the slope of the year is increasing (i.e. positive
## slope), and is randomly sampled between 0 and 0.4)
##
## - Situation 2: the slope is either increasing or decreasing, and is
## - sampled between -0.4 and 0.4.
lirs <- list(c(0,0.4),
             c(-0.4,0.4))
##
## Four trapping pressures are defined here (mean number of trapped
## animals per commune)
trap <- c(0.5, 1, 3, 10)

## Two intercepts for the model (either low prevalence = -3.1 or high
## prevalence -1.38).
situ <- c(-3.1, -1.38)
```

We then use the function `simulateIndicator()` to simulate each combination of the trapping pressure and epidemiological situation 1000 times and calculate both the true APC/mean prevalence levels and the indicators derived from the linear regression (see the help page of `simulateIndicator()`. WARNING!!!! THIS CALCULATION TAKES A VERY LONG TIME (several hours) !!!! Note that we have included the results of this calculation as a dataset of the package, so that the reader does not need to launch this function to reproduce further calculations:

```r
k <- 1
res <- list()
for (i in 1:4) {
    cat("** Trapping pressure:", trap[i],"\n\n")
    for (s in 1:2) {
        cat("Situation:", s,"\n\n")
        sin <- simulateIndicator(trap[i], cfcdo, situ[s],
                                 rangeSlope=lirs[[s]],
                                 nsim=1000, verbose=TRUE)
        sin$Situation <- c("Low Increasing","High")[s]
        sin$TrapPress <- paste0("mu = ",trap[i])
        res[[k]] <- sin
        k <- k+1
    }
}
resdfsim1 <- do.call(rbind, res)
resdfsim1$TrapPress <- factor(resdfsim1$TrapPress,
                              levels=c("mu = 0.5", "mu = 1",
                                       "mu = 3", "mu = 10"))
```

We load the results of these simulations:

```r
data(resdfsim1)
head(resdfsim1)

##    meanPrev           APC InterceptReg     SlopeReg
## 1 0.1167903  6.955361e-03    0.1127797 0.0093076703
## 2 0.1606609 -5.047723e-05    0.1697752 0.0001887324
## 3 0.1033724  4.691865e-03    0.1073033 0.0042334808
## 4 0.1072737  7.510773e-03    0.1105327 0.0136719452
## 5 0.1290129  1.420121e-02    0.1623073 0.0006260483
## 6 0.1887097  2.905227e-02    0.1979812 0.0309028260
```

```
##   SE_InterceptReg SE_SlopeReg Nind Ninfect      Situation
## 1      0.01473646 0.007517022  759     217 Low Increasing
## 2      0.01946637 0.009639504  640     187 Low Increasing
## 3      0.01485055 0.007690426  712     214 Low Increasing
## 4      0.01388698 0.006929551  807     222 Low Increasing
## 5      0.01867643 0.009512415  682     206 Low Increasing
## 6      0.01859711 0.009467817  769     229 Low Increasing
##   TrapPress
## 1  mu = 0.5
## 2  mu = 0.5
## 3  mu = 0.5
## 4  mu = 0.5
## 5  mu = 0.5
## 6  mu = 0.5
```

And we can summarize the results by plotting them (corresponding to Fig. 2 of the paper):

```
library(ggplot2)
ggplot(resdfsim1, aes(x=APC, y=SlopeReg))+
    geom_point(aes(col=TrapPress), alpha=0.5)+
    geom_abline(slope=1,intercept=0, lwd=1)+
    facet_grid(Situation~TrapPress)+coord_fixed()+
    xlab("True prop. of animals becoming infected in one year")+
    ylab("Estimated value by the regression")+
    theme(legend.position = "none")
```

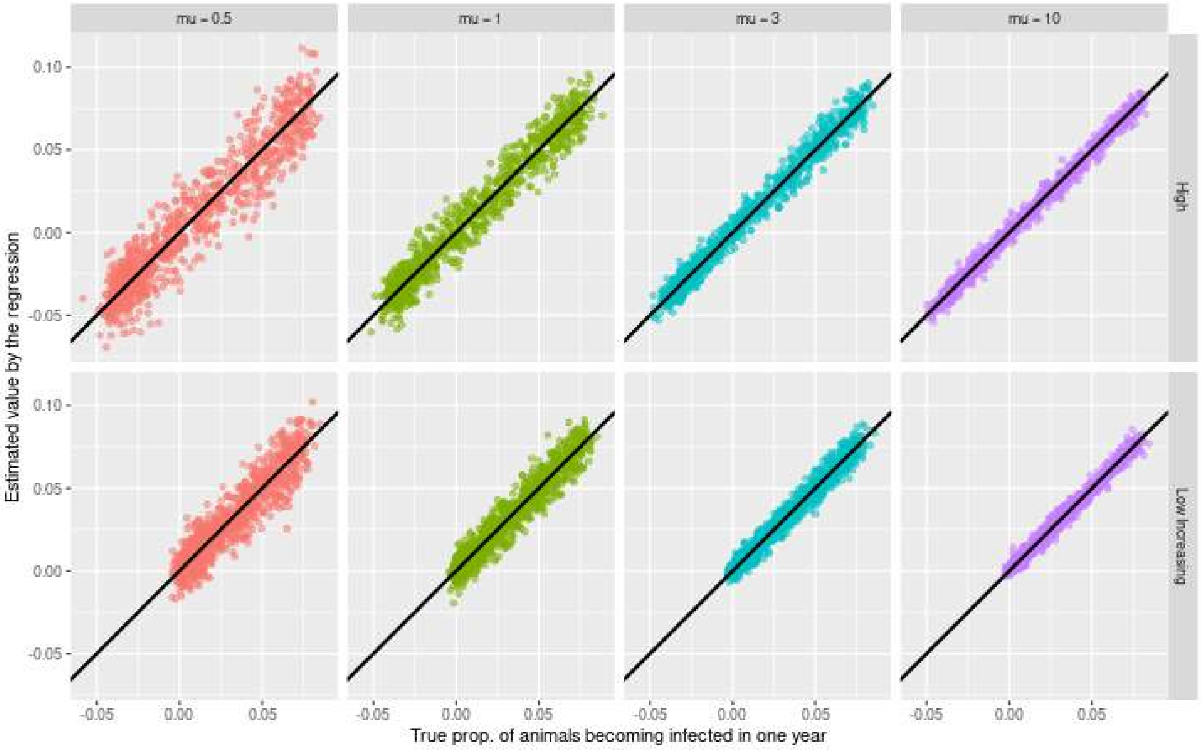

As noted in the paper, there is a good agreement between the two values. Note that even if these simulations focus on the ability of the regression model to recover the true value of the APC of the year, we can also examinate the ability of the intercept of the regression to recover the true mean prevalence level. Thus, even if we do not present these results in the paper, we can plot the mean prevalence estimated with the regression as a function of the true mean prevalence:

```
ggplot(resdfsim1, aes(x=meanPrev, y=InterceptReg))+
    geom_abline(slope=1,intercept=0, lwd=1, col="red")+
    geom_point(alpha=0.5)+
    facet_grid(Situation~TrapPress)+coord_fixed()+
    xlab("True mean Prevalence")+
    ylab("Estimated value by the regression")
```

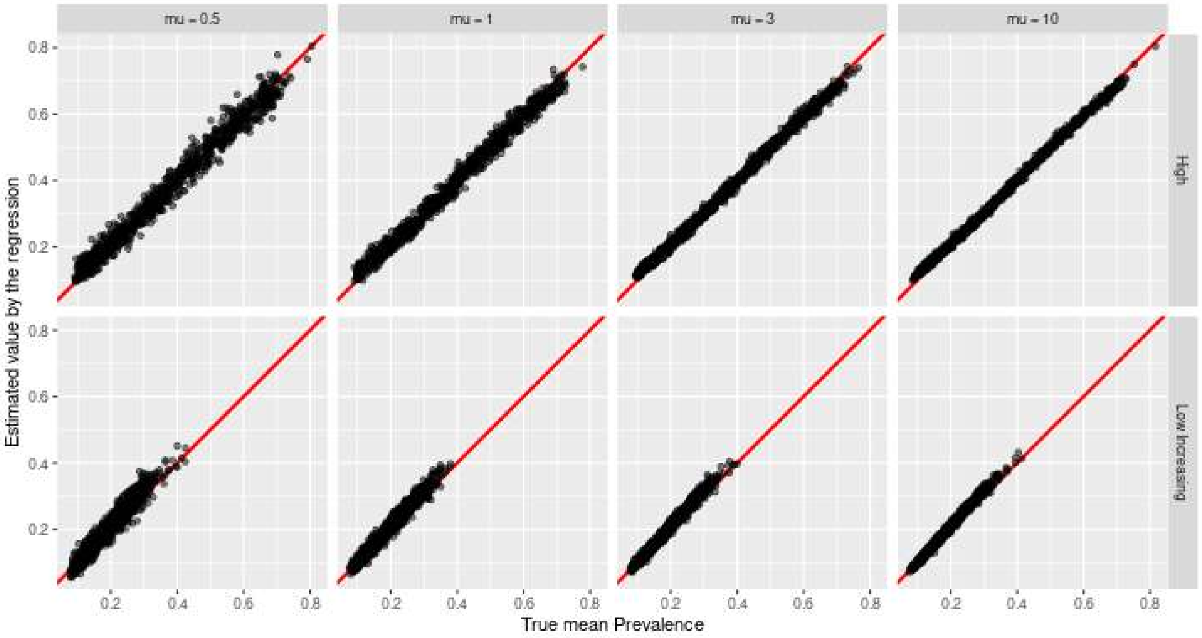

There is a good agreement between the true value and the estimated value. Finally, we can calculate the coverage probability of the 95% confidence intervals with the function `coverageCI()`:

```
cocia <- coverageCI(resdfsim1$APC,
                    resdfsim1$SlopeReg,
                    resdfsim1$SE_SlopeReg,
                    resdfsim1$Nind,
                    list(resdfsim1$TrapPress,
                         resdfsim1$Situation))
cocia
```

```
##        Situation TrapPress Probability
## 1           High  mu = 0.5       0.927
## 2           High    mu = 1       0.936
## 3           High    mu = 3       0.913
## 4           High   mu = 10       0.837
## 5 Low Increasing  mu = 0.5       0.942
## 6 Low Increasing    mu = 1       0.937
## 7 Low Increasing    mu = 3       0.893
```

```
## 8 Low Increasing   mu = 10       0.829
```

Which corresponds to the results provided Tab. 2 of the paper.

##### D.2.2 Second set of simulations

In the second set of simulations, we wanted to assess the ability of our regression model to estimate the mean prevalence level during the middle year of the study period. We simulated data with our Bayesian model, using different values of the intercept $\alpha = -4, -3, -2, -1, 0$, describing different mean prevalence levels. We then randomly sampled a slope $\beta$ in a uniform distribution bounded between -0.4 and 0.4. First, we define the parameters of the simulations:

```
## The intercepts
inter <- -c(4:0)

## The simulated trapping pressures
trap <- c(0.5, 1, 3, 10)
```

We then use the function `simulateIndicator()` to perform this second set of simulations. WARNING!!!! THIS CALCULATION TAKES A VERY LONG TIME (several hours) !!!! Note that we have included the results of this calculation as a dataset of the package, so that the reader does not need to launch this function to reproduce further calculations:

```
k <- 1
for (s in 1:4) {
    for (i in 1:5) {
        cat("** Prevalence level:", inter[i],"\n\n")
        sin <- simulateIndicator(trap[s], cfcdo, inter[i],
                                  rangeSlope=c(-0.4,0.4),
                                  nsim=1000, verbose=TRUE)
        sin$TrueIntercept <- inter[i]
        sin$TrapPress <- trap[s]
        res[[k]] <- sin
        k <- k+1
    }
}
resdfsim2 <- do.call(rbind, res)
resdfsim2$TrapPress <- paste0("mu = ", resdfsim2$TrapPress)
resdfsim2$TrapPress <- factor(resdfsim2$TrapPress,
                              levels=c("mu = 0.5", "mu = 1",
                                       "mu = 3", "mu = 10"))
resdfsim2$TrueIntercept <- paste0("alpha = ", resdfsim2$TrueIntercept)
resdfsim2$TrueIntercept <- factor(resdfsim2$TrueIntercept,
                                  levels=c("alpha = -4", "alpha = -3",
                                           "alpha = -2", "alpha = -1",
                                           "alpha = 0"))
```

We load the result of these simulations:

```
data(resdfsim2)
head(resdfsim2)

##      meanPrev           APC InterceptReg     SlopeReg
## 1 0.038753545  0.0004698377   0.03604526 -0.003695416
## 2 0.007664601 -0.0048136850   0.00724122 -0.003267482
```

```
## 3 0.023701063 -0.0017782333   0.02324332 -0.003694842
## 4 0.024237497 -0.0057497602   0.01421532 -0.001626778
## 5 0.114969045  0.0320244586   0.11600420  0.034870433
## 6 0.061259257  0.0100690054   0.05996670  0.008569524
##   SE_InterceptReg SE_SlopeReg Nind Ninfect TrueIntercept
## 1     0.008886886 0.004441970  754     223    alpha = -4
## 2     0.004182312 0.002049345  746     227    alpha = -4
## 3     0.007377678 0.003660030  696     212    alpha = -4
## 4     0.005805786 0.002967125  718     219    alpha = -4
## 5     0.013258408 0.006624248  835     248    alpha = -4
## 6     0.010305893 0.005361521  852     231    alpha = -4
##   TrapPress
## 1  mu = 0.5
## 2  mu = 0.5
## 3  mu = 0.5
## 4  mu = 0.5
## 5  mu = 0.5
## 6  mu = 0.5
```

We can now plot the results of the simulations, i.e. the mean prevalence estimated by the simple linear regressien vs the true mean prevalence:

```
ggplot(resdfsim2, aes(x=meanPrev, y=InterceptReg))+
    geom_abline(slope=1,intercept=0, lwd=1, col="red")+
    geom_point(alpha=0.5)+
    facet_grid(TrueIntercept~TrapPress)+coord_fixed()+
    xlab("True mean Prevalence")+
    ylab("Estimated value by the regression")
```

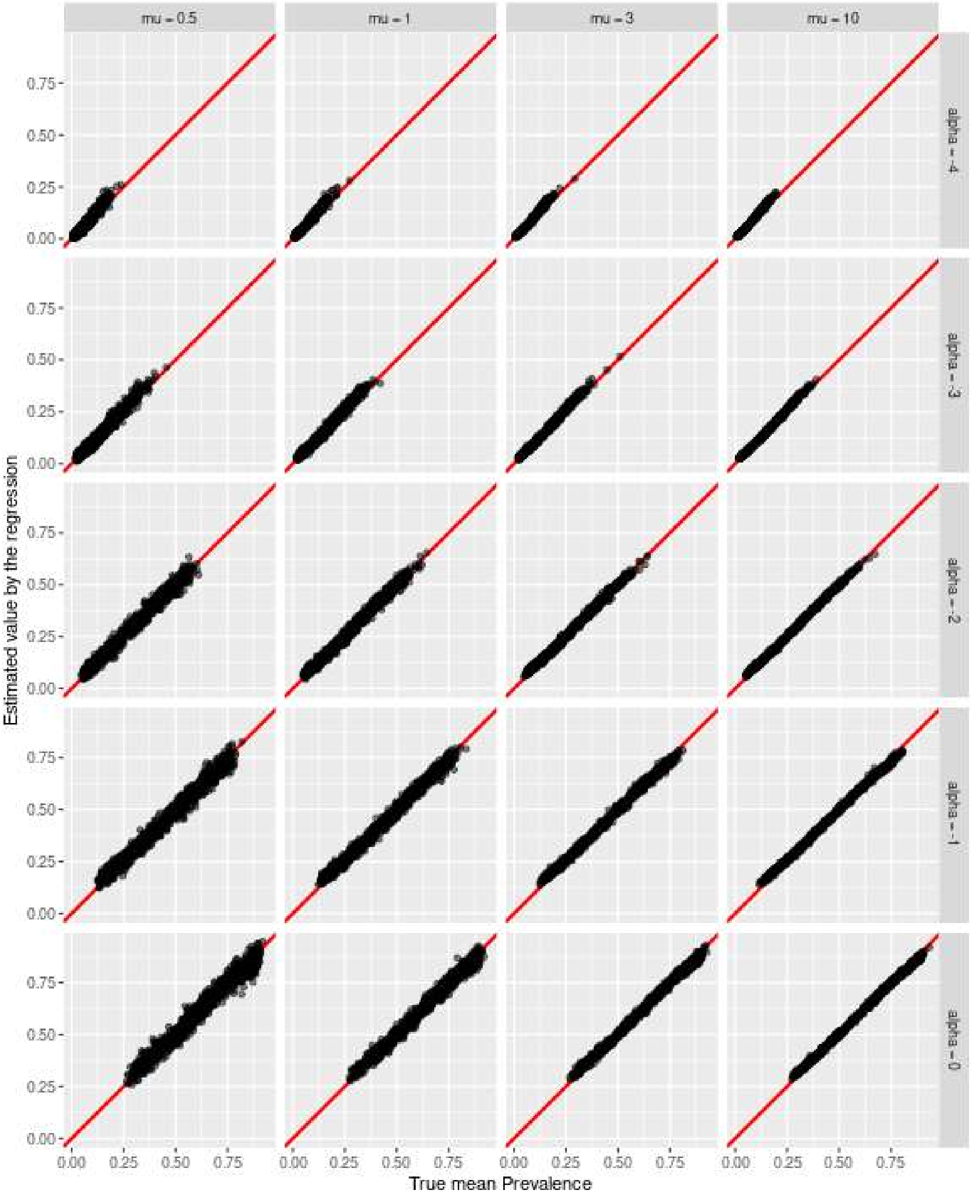

There is a close agreement between the two values. This figure corresponds to Fig. 3 of the paper.

Even if this set of simulations was designed to tests the ability of the regression intercept to estimate the true mean prevalence, we can still look at the ability of its slope to estimate the true APC, for our information (even if this plot is not shown in the paper):

```
ggplot(resdfsim2, aes(x=APC, y=SlopeReg))+
    geom_abline(slope=1,intercept=0, lwd=1, col="red")+
    geom_point(alpha=0.5)+
    facet_grid(TrueIntercept~TrapPress)+coord_fixed()+
    xlab("True prop. of animals becoming infected in one year")+
    ylab("Estimated value by the regression")
```

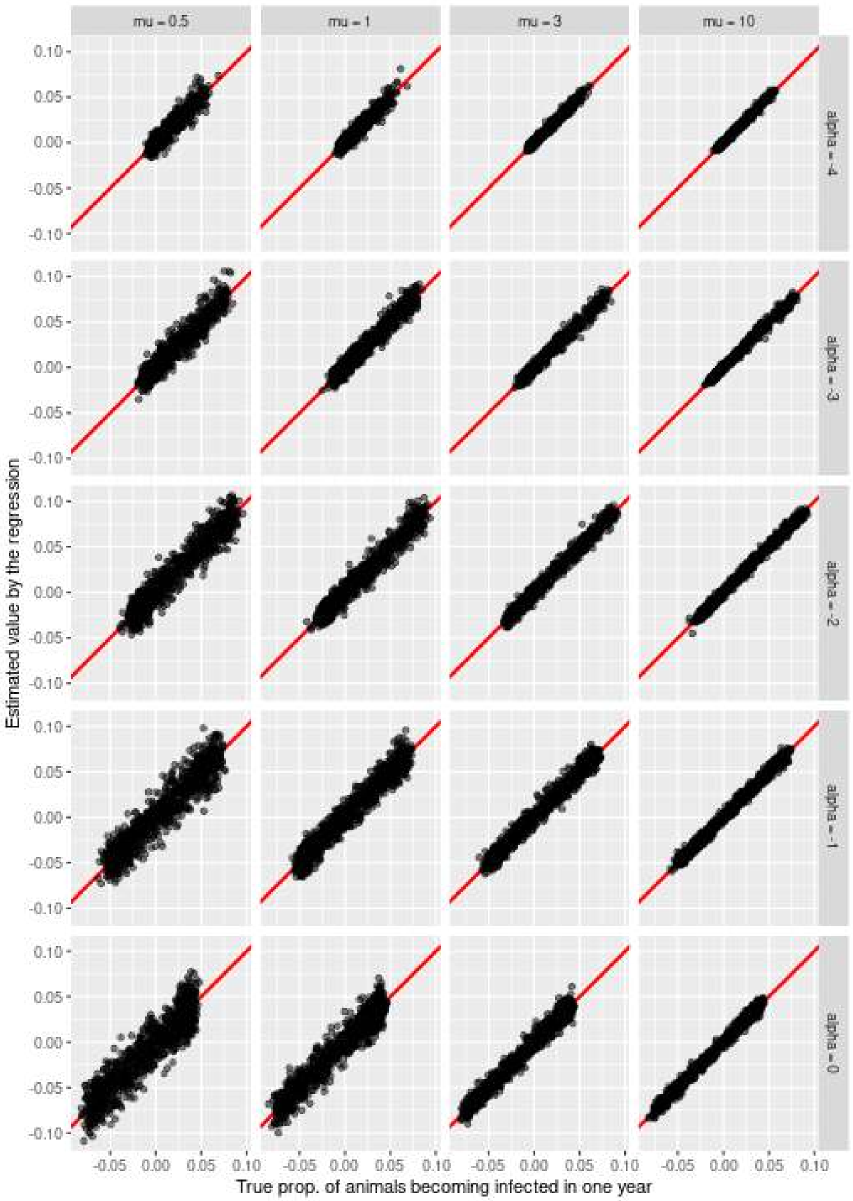

There is again a close agreement between the two values. Finally, we can use the function `coverageCI()` to estimate the coverage probability of the 95% confidence intervals of the coefficients:

```
cocim <- coverageCI(resdfsim2$meanPrev,
                    resdfsim2$InterceptReg,
                    resdfsim2$SE_InterceptReg,
                    resdfsim2$Nind,
                    list(resdfsim2$TrapPress,
                         resdfsim2$TrueIntercept))
names(cocim)[1] <- "Intercept"
cocim
```

```
##      Intercept TrapPress Probability
## 1  alpha = -4  mu = 0.5       0.912
## 2  alpha = -4    mu = 1       0.896
## 3  alpha = -4    mu = 3       0.781
## 4  alpha = -4   mu = 10       0.582
## 5  alpha = -3  mu = 0.5       0.935
## 6  alpha = -3    mu = 1       0.899
```

```
## 7  alpha = -3    mu = 3       0.785
## 8  alpha = -3   mu = 10       0.562
## 9  alpha = -2  mu = 0.5       0.931
## 10 alpha = -2    mu = 1       0.899
## 11 alpha = -2    mu = 3       0.803
## 12 alpha = -2   mu = 10       0.636
## 13 alpha = -1  mu = 0.5       0.921
## 14 alpha = -1    mu = 1       0.895
## 15 alpha = -1    mu = 3       0.771
## 16 alpha = -1   mu = 10       0.579
## 17  alpha = 0  mu = 0.5       0.917
## 18  alpha = 0    mu = 1       0.916
## 19  alpha = 0    mu = 3       0.865
## 20  alpha = 0   mu = 10       0.662
```

Which corresponds to Tab. 3 of the paper.

###### D.2.3 Third set of simulations

The last set of simulations was carried out to assess our regression model in more extreme situations. In all cases, we simulated data with our Bayesian model, using two different values of the intercept $\alpha = -2, 0$, describing different mean prevalence levels. We then randomly sampled a slope $\beta$ in a uniform distribution bounded between -0.4 and 0.4. We simulated two situations:

- We first simulated the spatial random effects with the iCAR model using a low value of $\tau = 0.1$ (instead of $\tau = 0.73$ in previous simulations) to simulate a very large spatial structure of the infection. When the spatial heterogeneity of the infection is large, there is a large remaining heterogeneity among the highly infected communes, which can pose a problem for the regression model.

- We then simulated a spatial structure of the infection changing with time. More precisely, we simulated two independent sets of commune random effects $\{u_i^{(1)}\}$ and $\{u_i^{(2)}\}$ using the iCAR model parameterized with $\tau = 0.73$. We then simulated data using a Bayesian model, using for each year $t$ the commune effects $\tilde{u}_i = ((t-1)/6) \times u_i^{(1)} + (1-(t-1)/6) \times u_i^{(2)}$, so that the commune effect transitions from the first set to the second set of commune effects. This simulation violated the assumption of additivity between space and time effects on the prevalence.

We simulated these two situations and calculated the two pairs of parameters of interest (mean prevalence and mean proportion of animals becoming infected in one year) under the two models (simulated Bayesian model and regression model), to assess the ability of the regression model to retrieve the true values. For each situation, we simulated two sampling designs: (i) the classical random sample used in previous situations, i.e. we generated a number of trapped animals for each year and each commune of the cluster from a binomial negative distribution with mean $\mu = 2$ and dispersion parameter $\theta = 0.48$; (ii) a strongly biased sample where one samples more individuals in more infected communes than in less infected communes, i.e. we generated a number of trapped animals for each year and each commune $i$ of the cluster from a binomial negative distribution with mean $\mu_i = 2 \times M \times u_i / (\sum_j u_j)$ (where $M$ is the total number of communes in the Dordogne/Charentes cluster) and dispersion parameter $\theta = 0.48$. Our aim was to demonstrate how biased sampling exacerbate the effect of the violation of hypotheses of additivity of space and time effects, and of small remaining spatial heterogeneity in highly infected communes.

To allow a point of comparison, we also simulated the same situation with random sampling, additivity of space and time effects, and moderate spatial heterogeneity (i.e. $\tau = 0.73$).

We use the function `simulateIndicator()` to perform this third set of simulations. WARNING!!!! THIS CALCULATION TAKES A VERY LONG TIME (several hours) !!!! Note that we have included the results of this calculation as a dataset of the package, so that the reader does not need to launch this function to reproduce further calculations:

```
res <- list()
interb <- c(0,-2)
k <- 1
for (b in 1:6) {
    for (i in 1:2) {
        cat("** Intercept:", i,"\n\n")

        sin <- simulateIndicator(2, cfcdo, interb[i],
                                 rangeSlope=c(-0.4,0.4),
                                 biasedSample=((b%%2)==0),
                                 nonadditive=(b>2&b<5),
                                 tau=c(0.1,0.73)[(b>2)+1],
                                 nsim=1000, verbose=TRUE)
        sin$TrueIntercept <- interb[i]
        sin$biasedSample <- ((b%%2)==0)
        sin$nonadditive <- (b>2&b<5)
        sin$tau <- c(0.1,0.73)[(b>2)+1]
        res[[k]] <- sin
        k <- k+1
    }
}

resdfsim3 <- do.call(rbind, res)
resdfsim3$Situation <- paste0(c("Additive","Interaction")[resdfsim3$nonadditive+1],", ",
                              c("moderate","strong")[(resdfsim3$tau<0.2)+1])
resdfsim3$Sampling <- c("random","directed")[resdfsim3$biasedSample+1]
resdfsim3$TrueIntercept <- paste0("alpha = ", resdfsim3$TrueIntercept)
resdfsim3$TrueIntercept <- factor(resdfsim3$TrueIntercept,
                                  levels=c("alpha = -2",
                                           "alpha = 0"))
```

We load the results of these simulations:

```
data(resdfsim3)
head(resdfsim3)

##    meanPrev         APC InterceptReg    SlopeReg
## 1 0.6310640 -0.05049301    0.6008090 -0.04540226
## 2 0.9210310  0.01445991    0.9121398  0.01345550
## 3 0.7084243 -0.01961578    0.7290296 -0.01561633
## 4 0.9112072  0.02299557    0.8991856  0.02974420
## 5 0.9378093  0.02694460    0.9271433  0.02825561
## 6 0.6455913 -0.03788675    0.6496327 -0.04315491
##   SE_InterceptReg SE_SlopeReg Nind Ninfect TrueIntercept
## 1      0.01349615 0.006760317 3454     233     alpha = 0
## 2      0.01494838 0.007373169 2818     213     alpha = 0
## 3      0.01438512 0.007394926 3089     233     alpha = 0
## 4      0.01334481 0.006403393 3500     224     alpha = 0
## 5      0.01416249 0.007178442 3123     220     alpha = 0
## 6      0.01431134 0.007253326 3178     236     alpha = 0
##   biasedSample nonadditive tau        Situation Sampling
## 1        FALSE       FALSE 0.1 Additive, strong   random
```

```
## 2          FALSE        FALSE 0.1 Additive, strong   random
## 3          FALSE        FALSE 0.1 Additive, strong   random
## 4          FALSE        FALSE 0.1 Additive, strong   random
## 5          FALSE        FALSE 0.1 Additive, strong   random
## 6          FALSE        FALSE 0.1 Additive, strong   random
```

And we can summarize the results by plotting them (corresponding to Fig. 5 We did not distinguish the two simulated intercepts $\alpha = 0$ and $\alpha = -2$ (we leave it to the reader to check that the conclusions are the same for the two values of $\alpha$ separately). First, we show the estimated mean proportion of animals becoming infected in one year:

```
ggplot(resdfsim3, aes(x=APC, y=SlopeReg))+
    geom_abline(slope=1,intercept=0, lwd=1, col="red")+
    geom_point(alpha=0.5)+
    facet_grid(Sampling~Situation)+coord_fixed()+
    xlab("APC")+
    ylab("Estimated slope by the regression")
```

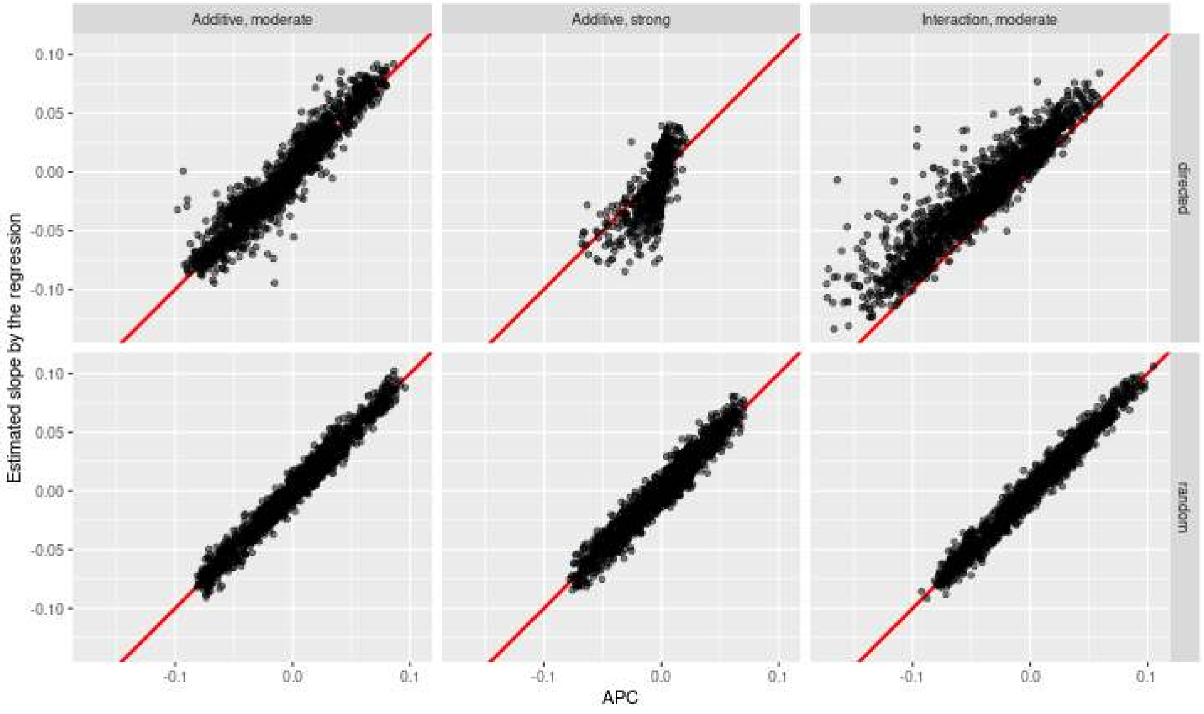

And then, the estimated mean prevalence of the infection:

```
ggplot(resdfsim3, aes(x=meanPrev, y=InterceptReg))+
    geom_abline(slope=1,intercept=0, lwd=1, col="red")+
    geom_point(alpha=0.5)+
    facet_grid(Sampling~Situation)+coord_fixed()+
    xlab("Mean prevalence")+
    ylab("Estimated intercept by the regression")
```

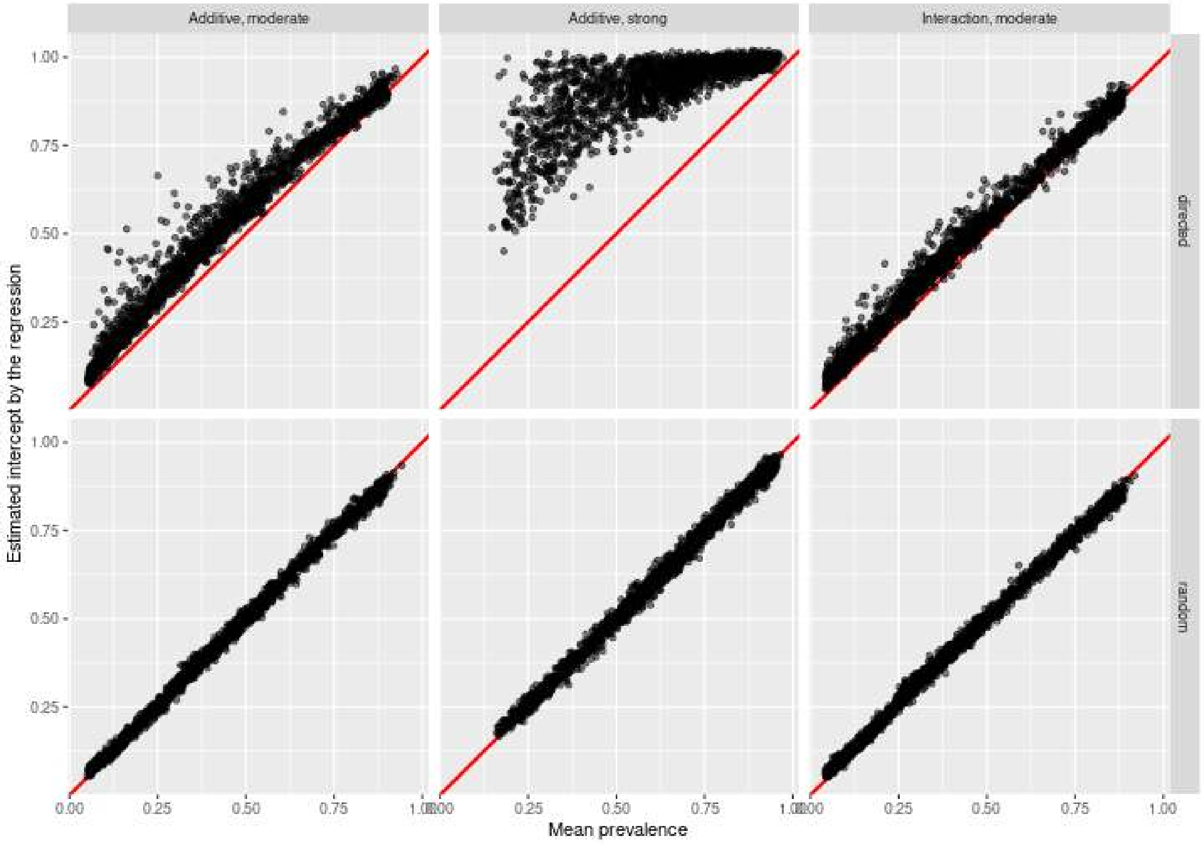

And finally, we can use the function `coverageCI()` to estimate the coverage probability of the 95% confidence intervals on the mean prevalence:

```
cocim3 <- coverageCI(resdfsim3$meanPrev,
                    resdfsim3$InterceptReg,
                    resdfsim3$SE_InterceptReg,
                    resdfsim3$Nind,
                    list(resdfsim3$Situation,
                         resdfsim3$Sampling))
names(cocim3)[1] <- "Intercept"
cocim3
```

```
##   Intercept             TrapPress Probability
## 1  directed    Additive, moderate      0.0665
## 2  directed      Additive, strong      0.0005
## 3  directed Interaction, moderate      0.2325
## 4    random    Additive, moderate      0.8680
## 5    random      Additive, strong      0.8815
## 6    random Interaction, moderate      0.7755
```

...and on the mean proportion of the population becoming infected in one year:

```
cocis3 <- coverageCI(resdfsim3$APC,
                    resdfsim3$SlopeReg,
                    resdfsim3$SE_SlopeReg,
                    resdfsim3$Nind,
                    list(resdfsim3$Situation,
                         resdfsim3$Sampling))
names(cocis3)[1] <- "Slope"
cocis3

##      Slope             TrapPress Probability
## 1 directed    Additive, moderate      0.6405
## 2 directed      Additive, strong      0.6850
## 3 directed Interaction, moderate      0.1900
## 4   random    Additive, moderate      0.9250
## 5   random      Additive, strong      0.9095
## 6   random Interaction, moderate      0.9200
```

So that generally, as long as the sample of trapped animals can be considered as a random sample, the model is robust to the violation of the underlying hypotheses. However, when the animals are preferentially trapped in the places where the prevalence is large, this exacerbate the bias in the parameters. Thus, the mean prevalence will be overestimated (and this bias will be larger when the spatial heterogeneity is strong), and the mean proportion of animals becoming infected in one year will also be biased (though this bias is much smaller than the one affecting the mean prevalence, and can be ignored for moderate spatial heterogeneity). Similarly, the nonrandom sampling can generate a bias in the estimation of the two parameters when both the spatial structure is changing with time and the sampling is directed towards the very infected communes.

### Appendix E Preliminary generalized additive models fitted to the datasets

#### E.1 Description of the modelling approach

As indicated in the main paper, we carried out an exploratory approach by fitting generalized additive models to the Sylvatub dataset, to check the assumption of additivity of space and time effects in the model.

Thus, we first fitted a generalized additive model (GAM, Wood, 2017) to explore the spatio-temporal structures of the infection. The commune is the finest available spatial resolution in our data, but we transformed our dataset into a spatial point pattern, to allow this preliminary exploration; we thus generated a random spatial location in the commune of trapping for each animal of the database. To account for the change of tests used to detect *M. bovis* infection after 2016, we subsampled the data after 2016: we randomly switched the infection status of one third of the animals tested positive after 2016 to “not infected”, thereby simulating a sensitivity of 50% after 2016. This ensured a constant sensitivity of the tests used to detect *M. bovis* during this first exploratory step (in the Bayesian model used in the paper, we take into account this variable sensitivity in a much better way). All these operations are programmed in the function `giveXYhomoSens()` of the package `badgertub` (see below).

We then fitted, for each French cluster of *M. bovis* infection, a binomial GAM predicting the infection status of the animals of the database (infected/not infected) as a function of space and time. Let $R_i$ denote the infection status due to *M. bovis* for the $i$th individual, $i = 1, ..., n$, which follows $R_i \sim \text{Bernoulli}(\pi_i)$, where $\pi_i$ denotes the probability of infection. A spatial-temporal GAM for modelling the infection risk with this dataset is formulated as follows:

$$\log\{\pi_i/(1-\pi_i)\} = \alpha_0 + f_t(\text{year}_i) + f_s(X_i, X_i) + f_{st}(X_i, X_i, \text{year}_i) \quad (6)$$

where $f_t(\text{year}_i)$ denotes an univariate thin plate spline (TPS) function of the year and $f_s(X_i, Y_i)$ denotes a bivariate TPS function of the spatial coordinates $X_i, Y_i$ of the animal $i$ (Wood, 2017). The interactive effect between space and time $f_{st}(X_i, Y_i, \text{year}_i)$ is a linear combination of tensor products between the bivariate TPS basis used for the spatial effect and the univariate TPS basis used for the time effect. More precisely, $f_{st}(\cdot)$ allows to focus on the interaction between the spatial and temporal dimensions, while marginal effects of space and time are accounted for by $f_t(\cdot)$ and $f_s(\cdot)$ (see Wood, 2017, p. 232). This allowed to test the significance of these spatio-temporal interactions, using the test developed by Wood (2017, section 6.2.1), derived from the approach of Nychka (1988).

We fitted these models using the package `mgcv` (Wood, 2017). Smoothing parameters for these smoothers were selected by REML, as this approach allows a the best approximation of P-values of significance tests of spatio-temporal interactions (Wood, 2017, p.262).

#### E.2 Implementation for the Dordogne/Charentes cluster

We first attribute coordinates to each trapped badger in the Dordogne/Charentes cluster, and subsample the data to homogenize the sensitivity of the tests:

```
set.seed(777)
xydo <- giveXYhomoSens(dotub$data, dotub$consts, cfcdo)
head(xydo)
```

```
##     y sensitivity year Nb ID        X       Y
## 1   0        0.50    2  3  1 428703.0 2053190
## 1.1 0        0.50    2  3  1 427720.8 2053897
## 1.2 0        0.50    2  3  1 426982.5 2053322
## 2   0        0.50    3  2  1 428740.2 2051209
## 2.1 0        0.50    3  2  1 426500.9 2053135
## 3   0        0.75    4  1  1 428928.6 2052880
```

We then fitted the full model with space-time interactions:

```
library(mgcv)
## Full model with space-time interactions and smoother for the year
gamdo_full <- gam(y~s(X,Y, k=40)+s(year,k=3)+
                      ti(X,Y,year, d=c(2,1), k=c(30,3)),
                  data=xydo, family=binomial, method="REML")
```

We then tested the significance of the space-time interactions:

```
summary(gamdo_full)
```

```
##
```

```
## Family: binomial
## Link function: logit
##
## Formula:
## y ~ s(X, Y, k = 40) + s(year, k = 3) + ti(X, Y, year, d = c(2,
##     1), k = c(30, 3))
##
## Parametric coefficients:
##             Estimate Std. Error z value Pr(>|z|)
## (Intercept)  -3.3932     0.1002  -33.86   <2e-16
##
## Approximate significance of smooth terms:
##                  edf Ref.df Chi.sq  p-value
## s(X,Y)        18.631 24.735 73.713 1.50e-06
## s(year)        1.000  1.001 17.302 3.28e-05
## ti(X,Y,year)   5.063  6.980  6.369    0.495
##
## R-sq.(adj) =  0.0293   Deviance explained = 8.62%
## -REML = 745.07  Scale est. = 1         n = 4323
```

These interactions are not significant. Note also that the estimated number of degree of freedom (`edf`) is equal to 1, confirming that the effect of the year is linear (as supposed by our Bayesian model).

#### E.3 Implementation for the Burgundy cluster

We then load the Sylvatub dataset for the Burgundy cluster, and then attribute coordinates to each trapped badger in this cluster, and finally subsample the data to homogenize the sensitivity of the tests:

```
data(botub)
data(cfcbo)

set.seed(777)
xybo <- giveXYhomoSens(botub$data, botub$consts, cfcbo)
```

We then fitted the full model with space-time interactions:

```
## Full model with space-time interactions and smoother for the year
gambo_full <- gam(y~s(X,Y, k=40)+s(year,k=3)+
                      ti(X,Y,year, d=c(2,1), k=c(30,3)),
                  data=xybo, family=binomial, method="REML")
```

We then tested the significance of the space-time interactions:

```
summary(gambo_full)

##
## Family: binomial
## Link function: logit
##
## Formula:
## y ~ s(X, Y, k = 40) + s(year, k = 3) + ti(X, Y, year, d = c(2,
##     1), k = c(30, 3))
##
## Parametric coefficients:
##             Estimate Std. Error z value Pr(>|z|)
```

```
## (Intercept)   -4.897      0.452  -10.84   <2e-16
##
## Approximate significance of smooth terms:
##                  edf Ref.df Chi.sq p-value
## s(X,Y)         20.23  25.02 51.356 0.00153
## s(year)         1.00   1.00  9.381 0.00219
## ti(X,Y,year) 12.80  17.64 22.032 0.23920
##
## R-sq.(adj) =  0.0975   Deviance explained = 24.1%
## -REML = 386.62  Scale est. = 1         n = 2786
```

Here again, these interactions are not significant. Note also that in this cluster too, the estimated number of degrees of freedom (`edf`) is equal to 1, confirming that the effect of the year is also linear here.

#### E.4 Implementation for the Bearn cluster

We finally load the Sylvatub dataset for the Bearn cluster, and then attribute coordinates to each trapped badger in this cluster, and finally subsample the data to homogenize the sensitivity of the tests:

```
data(betub)
data(cfcbe)

set.seed(777)
xybe <- giveXYhomoSens(betub$data, betub$consts, cfcbe)
```

We then fitted the full model with space-time interactions:

```
## Full model with space-time interactions and smoother for the year
gambe_full <- gam(y~s(X,Y, k=40)+s(year,k=3)+
                      ti(X,Y,year, d=c(2,1), k=c(30,3)),
                  data=xybe, family=binomial, method="REML")
```

We then tested the significance of the space-time interactions:

```
summary(gambe_full)

##
## Family: binomial
## Link function: logit
##
## Formula:
## y ~ s(X, Y, k = 40) + s(year, k = 3) + ti(X, Y, year, d = c(2,
##     1), k = c(30, 3))
##
## Parametric coefficients:
##             Estimate Std. Error z value Pr(>|z|)
## (Intercept)  -3.2617     0.1367  -23.86   <2e-16
##
## Approximate significance of smooth terms:
##                  edf Ref.df Chi.sq p-value
## s(X,Y)        12.312 17.056 24.298   0.112
## s(year)        1.001  1.001  1.259   0.262
## ti(X,Y,year)  8.154 11.745 17.891   0.110
##
## R-sq.(adj) =  0.0337   Deviance explained = 9.44%
## -REML = 374.01  Scale est. = 1         n = 1970
```

And here again, these interactions are not significant. Note also that in this cluster too, the estimated number of degrees of freedom (`edf`) is equal to 1, confirming that the effect of the year is also linear here.

#### References

Calenge, C. et al. in prep. – Assessing the dynamics of *Mycobacterium bovis* infection in three badger populations.

Gelman, A. and Meng, X. 1996. Model checking and model improvement. – In: Gilks, W. and Richardson, S. (eds.), Markov chain Monte Carlo in practice, chap. 11. Chapman & Hall/CRC, pp. 189–201.

Gelman, A. and Pardoe, I. 2007. 2. average predictive comparisons for models with nonlinearity, interactions, and variance components. – Sociological Methodology 37: 23–51.

Gelman, A. and Rubin, D. 1992. Inference from iterative simulation using multiple sequences. – Statistical Science 7: 457–472.

Gomila, R. 2020. Logistic or linear? estimating causal effects of experimental treatments on binary outcomes using regression analysis. – Journal of Experimental Psychology in press.

Hellevik, O. 2009. Linear versus logistic regression when the dependent variable is a dichotomy. – Quality & Quantity 43: 59–74.

Hisakado, M. et al. 2006. Correlated binomial models and correlation structures. – Journal of Physics A: Mathematical and General 39: 15365.

Martin, J. et al. 2011. Accounting for non-independent detection when estimating abundance of organisms with a bayesian approach. – Methods in Ecology and Evolution 2: 595–601.

Nychka, D. 1988. Bayesian confidence intervals for smoothing splines. – Journal of the American Statistical Association 83: 1134–1143.

Wood, S. N. 2017. Generalized Additive Models: An Introduction with R. Chapman & Hall/CRC Texts in Statistical Science, 2 edn. – Chapman and Hall / CRC.
